# A functional interleukin-4 homolog is encoded in the genome of infectious laryngotracheitis virus: unveiling a novel virulence factor

**DOI:** 10.1101/2024.07.10.602945

**Authors:** Jeremy D. Volkening, Stephen J. Spatz, Maricarmen García, Teresa A. Ross, Daniel A. Maekawa, Kenneth S. Rosenthal, Ana C. Zamora, April Skipper, Julia Blakey, Roshan Paudel

**Author notes:** Corresponding author *Email address:* (Stephen J. Spatz).

## Abstract

Herpesviruses have evolved numerous immune evasion tactics, persisting within their hosts through self-perpetuating strategies. One such tactic involves acquiring functional copies of host genes encoding cytokines such as IL-6 (HHV-8), IL-10 (HHV-4, HHV-5), and IL-17 (SaHV-2). These viral mimics, or virokines, can bind to cellular receptors, modulating the natural cytokine signaling to manipulate the immune response in favor of the virus or stimulate target cell growth to enhance virus replication. In the course of full-length cDNA sequencing of infectious laryngotracheitis virus (ILTV) transcripts, a previously unknown highly-spliced gene was discovered in the viral genome predicted to encode a 147 amino acid protein with similarity to vertebrate interleukin-4. The three-intron gene structure was precisely conserved with chicken and other vertebrate IL-4 homologs, and the amino acid sequence displayed structural conservation with vertebrate homologs at the primary, secondary, and tertiary levels based on computational modeling. The viral IL-4 gene was subsequently identified in all sequenced ILTV genomes. The mature transcript was highly expressed both *in vitro* and *in vivo*, and protein expression in infected cells was confirmed using LC-MS/MS. Phylogenetic analyses, along with the conserved gene structure, suggested direct capture from a *Galliformes* host. Functionally, an LPS-stimulation assay showed that the expressed viral IL-4 homolog stimulated nitric oxide production in a macrophage cell line at comparable levels to recombinant chicken IL-4. A recombinant virus lacking vIL-4 exhibited slightly higher titers in cell culture compared to the parental strain. *In vivo* bird studies demonstrated reduced pathogenicity of the vIL-4 knockout compared to wildtype. These results represent the first report of a previously unknown virokine encoded in the ILTV genome expressing a functional IL-4 homolog and virulence factor.

**Author Summary:** Herpesviruses are large DNA viruses, several of which express proteins that are homologous to host cytokines (termed virokines) and are thought to modulate the host immune response in favor of the virus. We report the identification of a novel virokine in the genome of infectious laryngotracheitis virus (ILTV) that is structurally and functionally similar to avian interleukin-4. The significance of this finding is threefold:

1. **Novel mechanism**: The identification of vIL-4 expands our understanding of the strategies employed by viruses to evade the host immune response, including through the acquisition, adaptation, and expression of cellular genes for their own advantage.
2. **Implications for disease pathogenesis**: We demonstrate that vIL-4 plays a functional role in ILTV virulence. Understanding the mechanisms by which this happens could lead to the development of novel therapeutic strategies.
3. **Evolutionary insights**: The presence of the highly spliced vIL-4 gene in the ILTV genome suggests direct genomic capture of a host gene, providing insights into the evolutionary history of ILTV and its interactions with avian theropods.

Overall, this research represents a significant contribution to our understanding of viral immunology and has potential implications for the development of improved vaccines for ILTV infection.

## Introduction

Infectious laryngotracheitis virus (ILTV) causes acute upper respiratory disease that impacts chickens reared on high-density poultry farms, with severity of the disease dependent on the viral strain, infectious dose, and age of the bird [1,2]. ILTV replicates to high titers in the laryngeal and tracheal mucosae, leading to severe cytolytic damage to mucosal epithelia and hemorrhaging. Mild disease causes coughing, nasal discharge, and conjunctivitis; severe disease is further characterized by marked dyspnea, gasping, open-mouth breathing, and expectoration of bloody mucoid phlegm [1]. Mucoid plugs, likely the result of mucus hypersecretion by goblet cells in the trachea, often obstruct airways, predisposing birds to death due to asphyxiation [3]. If birds survive acute infection, a lifelong latent infection of the trigeminal ganglia will remain.

As with other *α*-herpesviruses, it is well-documented that cell-mediated immunity is required for resolution of ILTV infection by the immune response. B cells and antibody are not essential for protection since chicks bursectomized within 24 hours post-hatch and vaccinated four weeks later are protected from ILTV challenge [2]. In addition, these results are consistent with previous findings from passive transfer experiments which showed that transferring antibodies from hyperimmune chickens does not provide protection against infection [4,5]. However, thymectomized chickens are very susceptible to disease as shown in challenge experiments [4,6]. Sabir et al. recently found only a weak association between protective immunity against ILTV and antibody titers to individual ILTV antigens such as glycoproteins C, D, E, G, I, and J [7]. These and other studies indicate a pivotal role for cell-mediated immunity in combating ILTV infection [8].

Viruses utilize their host for more than replication and have acquired host genes that support or facilitate their replication and parasitism of the host. Larger DNA viruses, which likely captured most of their non-core genes from host genomes during the course of viral evolution [9], have physical room for a relatively diverse gene repertoire encompassing specialized roles beyond replication and structure in immune invasion, pathogenicity, and other functions that confer an evolutionary advantage. Among these specialized host-acquired genes are those which manipulate the host immune response, including functional homologs of cytokines and their receptors. These genes provide agonistic or antagonistic effects relative to their host counterparts [10]. Within the herpesviruses, at least four viruses harbor host-acquired genes for cytokine homologs: IL-6 in HHV-8; IL-17 in SaHV-2; and IL-10 in both HHV-4 and HHV-5 [11–14]. These are termed virokines.

The dsDNA type D genome of ILTV is approximately 151 Kbp in length and encodes approximately 80 putative protein-coding open reading frames (ORFs). The majority of ORFs are homologous to genes of the *α*-herpesvirus prototype HSV-1 and also include eight ORFs clustering near the termini of the unique long subgenomic region (A, B, C, D, E, F, UL-0 and UL-1) that are unique to members of the *Iltovirus* genus. An earlier study classified the genes of ILTV into four categories based on their transcription kinetics: immediate early (*α*), early (β), leaky late (γ1) and true late (γ2) [15]. Immediate early genes are produced in the absence of *de novo* viral protein synthesis. Early viral genes typically code for enzymes necessary for DNA replication, while late genes encode for the structural polypeptides of the virion and additional proteins to facilitate infection. Transcription in ILTV is largely “leaky” (not strictly temporally controlled), with only ICP4 expression qualifying as truly immediate early. This is similar to equine herpesvirus 1 (EHV-1) which also encodes a single *α* gene (ORF62, an ICP4 homolog) [16]. In contrast, HSV-1 expresses four immediate-early genes (ICP4, ICP0, ICP22, and ICP27) [17].

Studies investigating the transcriptome of ILTV-infected cells have employed both *in vitro* and *in vivo* models and have relied on short-read sequencing (SRS) for assembling the viral and cellular transcripts [18–21]. Although SRS has a high yield and base accuracy, its short read length is ill-suited to identifying alternate splice forms and overlapping/co-terminal transcripts such as are common in herpesviruses [22]. Conversely, long-read sequencing (LRS) technologies such as Oxford Nanopore (ONT) and Pacific Biosciences (PacBio) are capable of resolving longer, in many cases full-length, transcript isoforms with varying rates of accuracy. LRS can be used to redefine the transcriptomes of virus-infected cells given sufficient viral coverage, sometimes in combination with deep SRS sequencing [23–25].

In the course of an effort to define the viral transcriptome of primary chicken kidney cells infected with either wild type (1874C5) ILTV, a US clade VI strain, or the vaccine CEO strain Laryngo-Vac (Zoetis) using PacBio IsoSeq full-length cDNA sequencing, we discovered a novel spliced transcript family mapping to a previously unannotated genomic region at the UL/IRS junction and predicted to encode a homolog to avian interleukin 4. We hypothesize that ILTV acquired an ancient avian interleukin 4 gene through a non-retrotranscription (direct) capture mechanism during host coevolution which potentially aided immune evasion or viral replication. We demonstrate that vIL-4 protein is expressed, exhibits structural and functional properties similar to those of chicken IL-4, and can modify macrophage function. To investigate the role of vIL-4 during ILTV infection, a deletion mutant virus lacking vIL-4 (ΔvIL-4) was generated and characterized in terms of its growth kinetics in cell culture and pathogenicity in birds.

## Results

### ILTV encodes a novel spliced transcript at the U_S_/IR_S_ junction

A novel transcript family mapping to a previously unannotated region of the ILTV genome was revealed in the course of full-length cDNA sequencing of the ILTV transcriptome. The primary isoform consists of four exons transcribed from the end of the IRS, on the opposite strand to the surrounding UL(−1) and ICP4 genes (Figure 1). This isoform was detected in the IsoSeq transcripts from both virulent (1874C5) and attenuated (CEO Laryngo-Vac) strains. Less abundant junction-spanning splice forms were also observed in 1874C5, although at <1% of the abundance of the primary transcript. Post-discovery, we were also able to detect the transcript in existing short-read data from the NCBI Sequence Read Archive (an example is shown in Figure 1).

**Figure 1:**
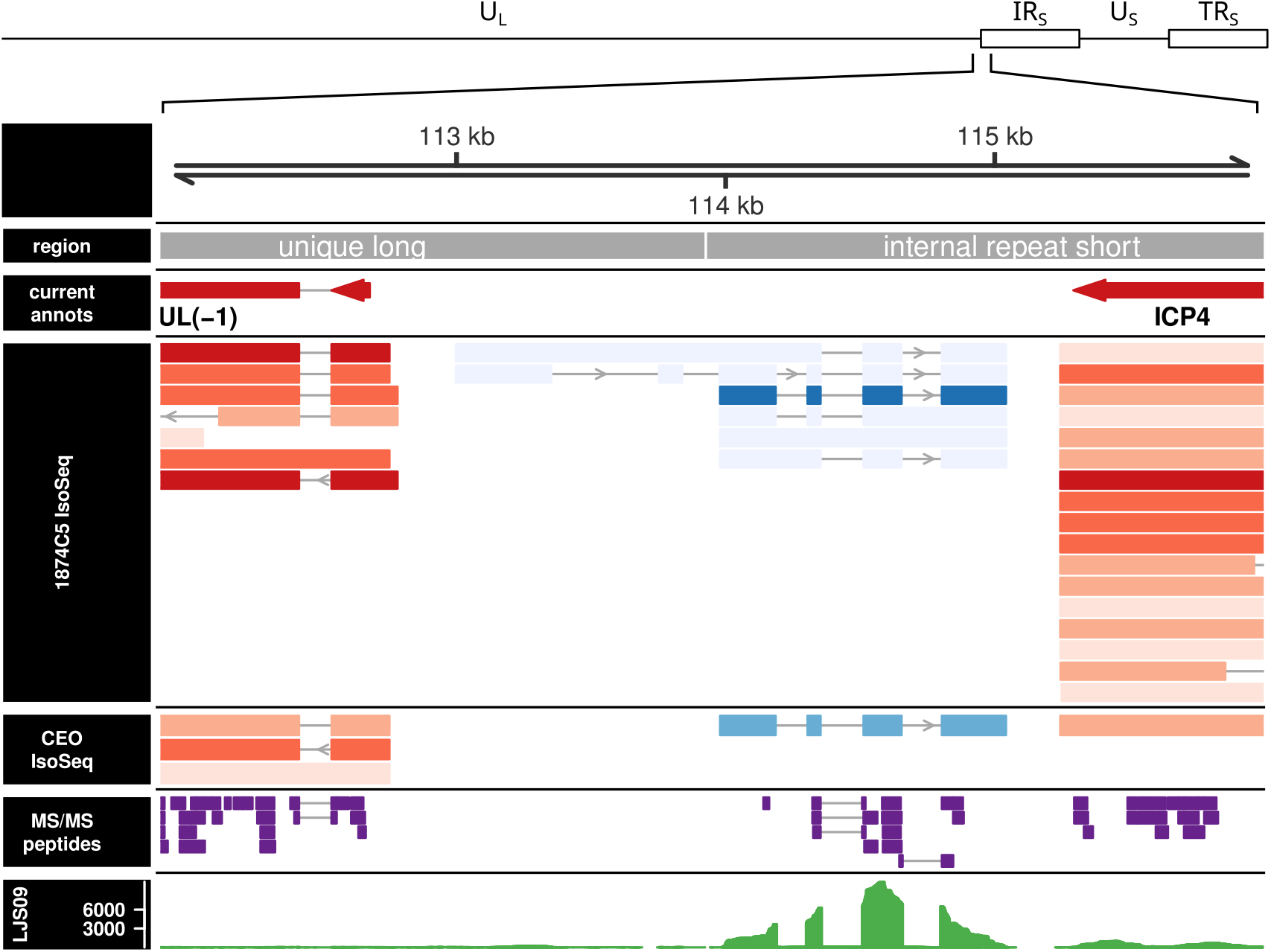
Alignment of experimental IsoSeq transcripts and MS/MS peptides to the ILTV UL/IRS genomic junction. Tracks shown are (from top to bottom) genomic coordinates; genomic regions; existing annotations; 1874C5 IsoSeq models; CEO (LaryngoVac) IsoSeq models; LC-MS/MS experimentally identified peptides; published LJS09 RNA-Seq read depth. IsoSeq models are colored by strand (blue: +, red: −) and shaded by supporting read count (darker = higher count). The primary 4-exon vIL-4 transcript (center, forward strand) is roughly 100× more abundant than the alternative isoform models in 1874C5. In CEO (LaryngoVac), where there was lower viral abundance overall, only the dominant isoform was detected. Most of the theoretically detectable tryptic peptides from vIL-4 were detected at 1% FDR using bottom-up LC-MS/MS (purple features). Also shown (bottom track) is an alignment of previously published short-read RNA-Seq data from strain LJS09 demonstrating expression of the vIL-4 transcript in an additional strain. Only a single replicate from this experiment is shown for the sake of brevity, but all four infected replicates showed similar coverage.

The novel transcript (named *vIL-4* as discussed below) was found to be highly abundant both *in vitro* and *in vivo*. In the *in vitro* semi-quantitative IsoSeq data, the *vIL-4* transcript was present at 3247 and 2606 viral transcripts per million (vTPM) in 1874C5 and CEO/LaryngoVac, respectively. By rank, these equated to the 18th and 19th most abundant transcript in each strain. Short-read RNA-Seq revealed the *vIL-4* transcript to be even more abundant *in vivo*, representing the fifth-most abundant viral transcript in birds inoculated with either 1874C5 or CEO/LaryngoVac (Figure 2).

**Figure 2:**
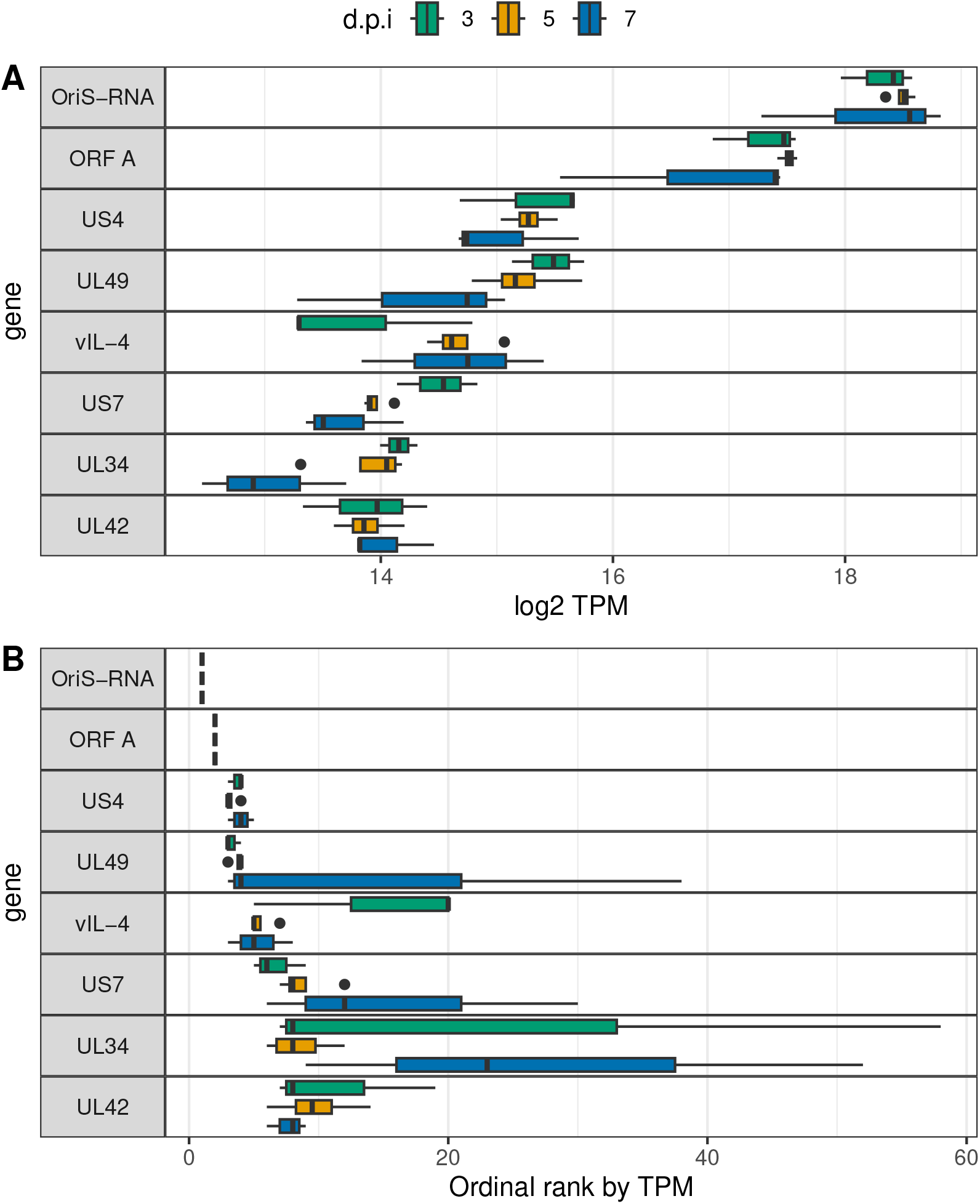
Highly expressed in vivo viral transcripts. Shown are the 8 most highly expressed ILTV transcripts by median transcripts per million transcripts (TPM) as a distribution of (A) viral TPM and (B) ordinal rank of TPM. RNA-Seq expression data is from ocular innoculation of birds with 1874C5 followed by sampling at 3, 5, and 7 days post-infection (d.p.i.). In this experiment, vIL-4 is the fifth most abundant transcript expressed, following two origin-associated transcripts, glycoprotein G, and VP22.

### The novel transcript codes for an IL-4 homolog

The longest open reading frame of the novel transcript codes for a 147 amino acid protein. A BLASTp search of the *nr* database suggested closest similarity to avian interleukin-4 homologs; the gene has thus been named *vIL-4* in keeping with nomenclature from other herpesvirus virokines. In order to carefully examine the relationship between vIL-4 and vertebrate homologs, the IL-4 genes from 29 sequenced avian and related crocodilian and turtle genomes were carefully curated. Species were chosen to broadly represent the avian tree of life, with the requirement that the full IL-4 gene structure was locatable in the genome. Some gene structures already existed in the genome annotations; in other cases, gene structures were determined by a combination of protein-to-nucleotide alignment and *de novo* gene prediction as described in the methods.

Secondary and tertiary structure prediction showed vIL-4 to be a classic four-helical cytokine with high confidence (Figures 3 and 4). It contains the expected N-terminal signal peptide and four universally conserved cysteine residues forming stabilizing disulfide bridges across the fourhelical bundle. The predicted tertiary structures of vIL-4 and chicken IL-4 bound to the Type I receptor complex aligned closely (Figure 4C). In fact, spatial analysis of the predicted models suggests a potentially stronger binding interaction between vIL-4 and its putative type I receptor IL-4Rα and γC (CD132) subunits than that of cIL-4 (21 and 17 predicted hydrogen bonds+salt bridges, respectively)(Figure 7F).

**Figure 3:**
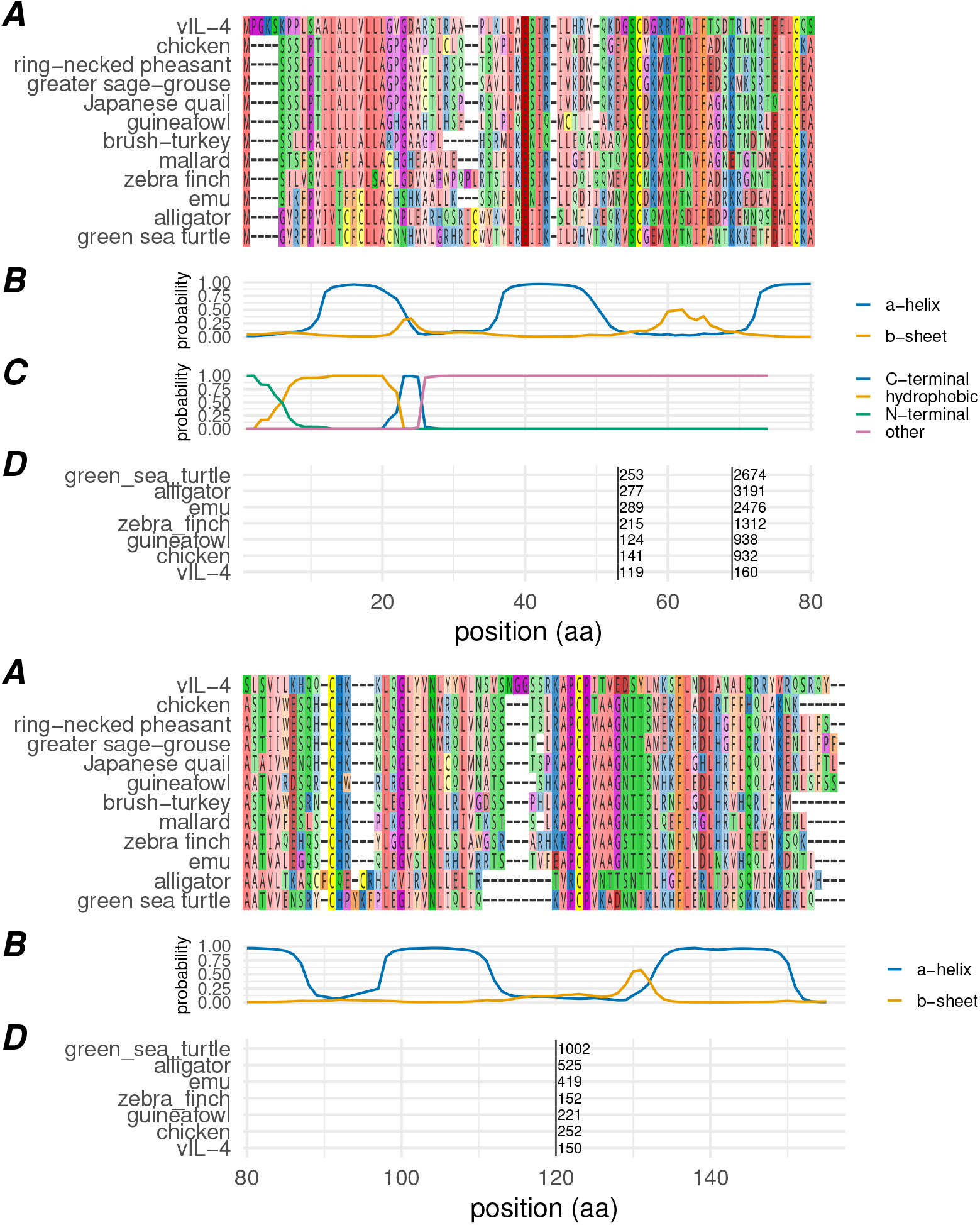
Primary, secondary, and genomic structure of the viral interleukin-4 homolog. (A) Amino acid alignment of vIL-4 with homologs from selected species. (B) Secondary structure probability of the vIL-4 amino acid sequence as predicted by JPRED2. (C) Signal peptide domain probability as predicted by SignalP 6.0; indicated are the N-terminal domain [‘N’], hydrophobic domain [‘H’], and C-terminal domain [‘C’]. (D) Location of genomic introns in a representative selection of homologs, translated into amino acid space. Black vertical bars indicate the intron position, with the intron length indicated to the right of each bar. Intron locations within the amino acid alignment were identical for all homologs examined, including those not shown.

**Figure 4:**
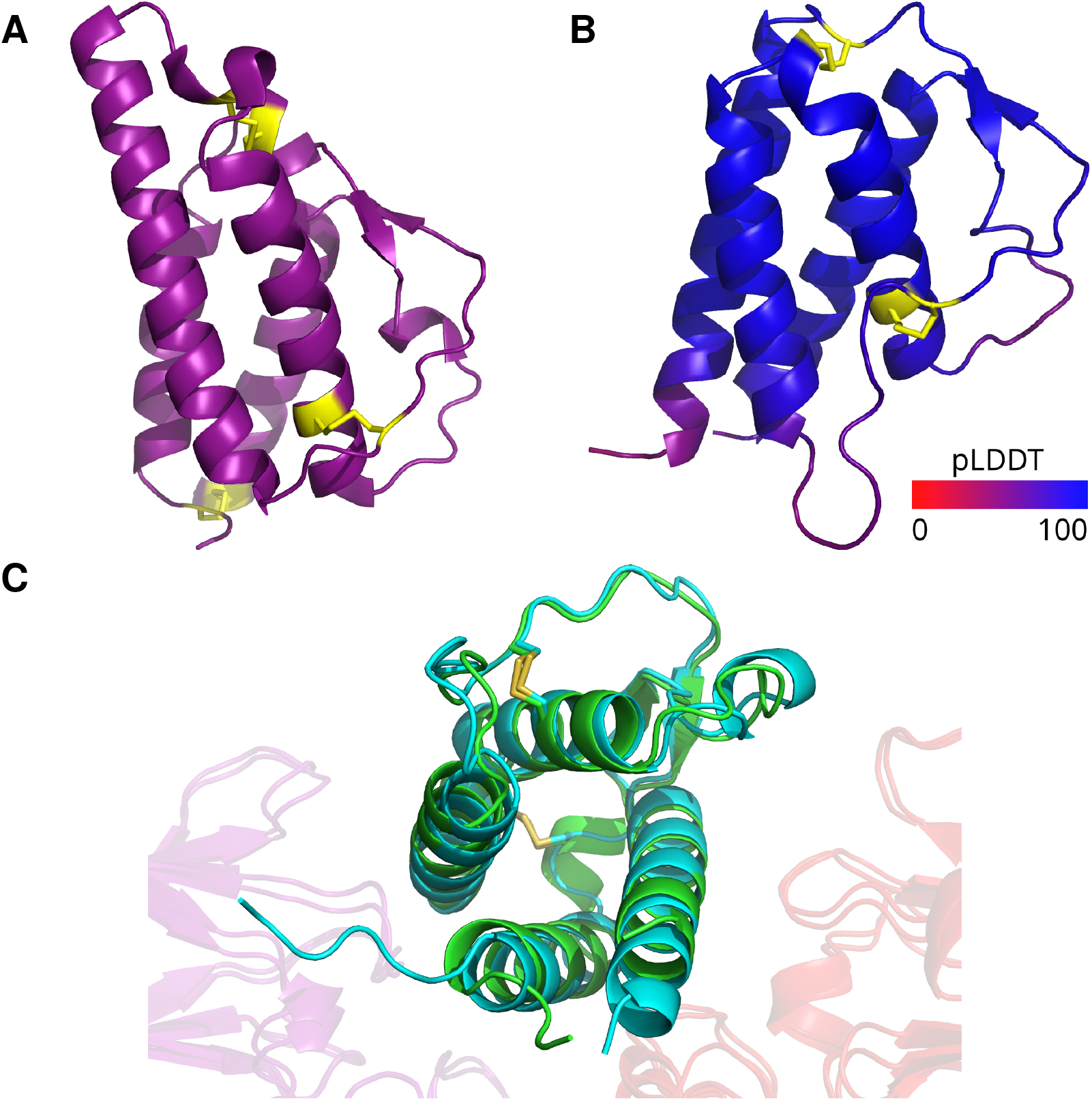
Comparative predicted tertiary structures. (A) Empirical crystal structure of human IL-4 (PDB:2B8U); cysteines/disulfide bonds shaded yellow. (B) AlphaFold model of vIL-4 shaded by pLDDT from low (red) to high (blue); cysteines/disulfide bonds shaded yellow. ILTV vIL-4 is predicted to share the same four-alpha-helix, anti-parallel beta sheet arrangement as the empirically modeled hIL-4. Like all avian IL-4 homologs, it lacks the cysteines at both protein termini that form a third disulfide bond in human IL-4 (panel A lower-left). (C) Superimposed predicted structures of vIL-4 (blue) and chicken IL-4 (green) in complex with chicken IL-4R α (transparent purple) and γc (transparent red) receptor subunits (Type I receptor complex), as predicted by AlphaFold and aligned by the RCSB Pairwise Structure Alignment tool (jFATCAT/rigid). Disulfide bridges are shown in yellow.

The genomic structures of the 30 curated IL-4 genes, including *vIL-4*, all contained three introns with identical locations relative to the amino acid alignment, and all intron boundaries occurred without exception at phase 0 in the coding sequence. The introns were located between core secondary structures (the first α-helix, first β-strand, α-helices two and three, and α-helix four) (Figure 3). All three of these characteristics (three coding sequence introns, aligned to codon phase 0, and occurring between adjacent secondary structures) have previously been shown to be conserved within the four-helical cytokine family as a whole [26]. Intron lengths varied widely, generally increasing with increasing genetic distance from vIL-4. Conservation of the vertebrate genomic structure in vIL-4 is in contrast to other herpesvirus-encoded cytokines, which are either unspliced or contain a single intron.

### Viral IL-4 protein is expressed in cell culture

Bottom-up proteomics of total protein from ILTV-infected cell culture detected six fully trypic peptides and numerous missed-cleavage peptides from vIL-4 at a 1% peptide FDR (Figure 1; Table S2). No peptides were detected from within the predicted N-terminal leader sequence. If that region is ignored, six of eight possible trypic peptides (within the detectable size range of the method used) were detected, covering 86% of the detectable sequence (Figure S5). This includes peptides spanning two of the three intron junctions. Identification QC plots for the primary proteoforms of each detected peptide are shown in Figures S6-S18. All identified peptides were represented by MS2 spectra with rich y/b ion series and all but one had precursor extracted ion chromatograms specific only to infected cells. The exception, peptide LNETEELCQSLSVILK, had a bimodal elution profile at the target m/z and retention time window, one peak of which was shared by the mock-infected samples and the other not (Figure S11); it is therefore inconclusive as support for that specific peptide identification. Overall, the identified peptides represent high-quality spectral matches and provide clear evidence for translation of vIL-4 protein specific to infected cells.

Based on label-free peptide quantification with FlashLFQ and protein-level summarization using the riBAQ metric, vIL-4 comprised 0.94% of all viral protein by molarity in cells infected with the LaryngoVac vaccine strain, and 0.70% of viral protein in cells infected with the group VI virulent strain 1874C5. This placed it as the 22nd and 23rd most abundant protein out of 66 and 64 viral proteins detected, respectively.

### *Viral IL-4 evolved within* Galliformes

Viral IL-4 had a global amino acid identity to chicken IL-4 of 34.7% (max: turkey, 39.5%) and a global similarity to chicken IL-4 of 52.4% (max: Japanese_quail, 56.8%). Multiple alignment within this representative group (Figure 5). Trees produced from amino acid and nucleotide coding sequence alignments were highly congruent.

**Figure 5:**
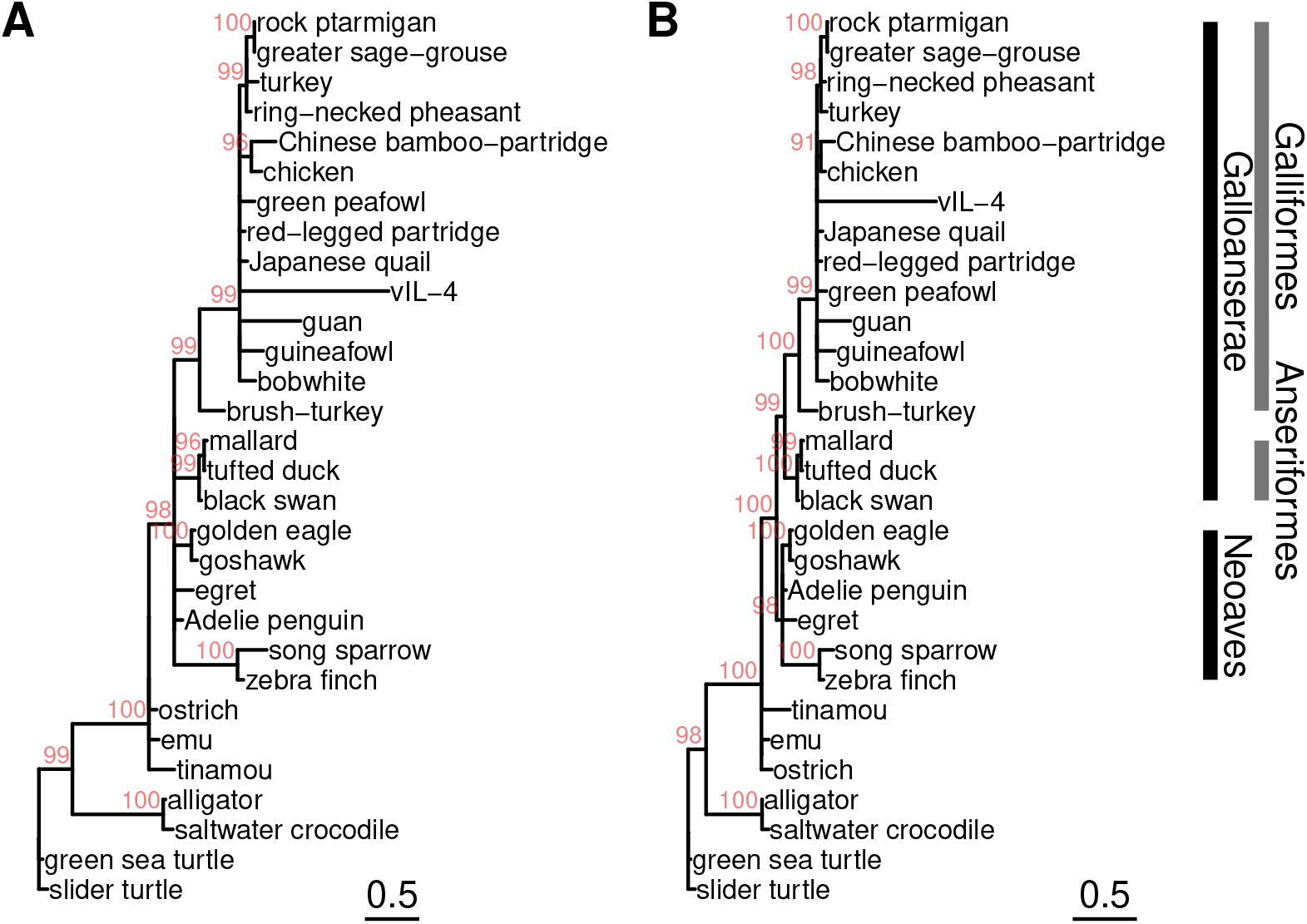
ML trees of vIL-4 and interleukin-4 homologs from a selection of representative bird and reptilian genomes. (A) From amino acid alignment, built with IQTREE using the ‘Q.bird+I+G4’ model as selected by ModelFinder, ultrafast bootstrapping (1000 iterations). (B) From coding nucleotide alignment, built with IQTREE using the ‘GTR+I+R’ model as selected by ModelFinder, ultrafast bootstrapping (1000 iterations). Branches with UFBoot support (in red) < 90 were collapsed.

Viral IL-4 did not partition most closely with chicken, but rather appears to have emerged from a more distant common clade representing the Galliformes subsequent to divergence of brush-turkey (Fig. 5). Overall, the phylogenetic distance within this group was small. Based on the analysis of Chen et al. [27], this clade diverged roughly 75 million years ago (Mya) in the Late Cretaceous Epoch. There are several possible interpretations of this result. First, ILTV may have captured the host IL-4 from a distant ancestor during this period of divergence. This would place the origins of ILTV vIL-4 much further in the past than current estimates of divergence between sequenced present-day strains, which have been estimated at only 1–1.5 Mya [28], but on a similar time scale to the acquisition of a host PD-1 receptor ligand by TTGHV1 roughly 50 Mya [29]. Another possibility is that vIL-4 was captured more recently from an as-of-yet unsequenced avian host. While ILTV is most commonly associated with chickens, it has also been reported to infect pheasant, partridges and peafowl [1,30]. It is also possible that the different rates of evolution between viral and vertebrate homologs confounds the ability to correctly interpret the phylogenetic placement. The IL-4 trees suggest that vIL-4 is evolving at a rate at least twice as fast as the next most divergent sequence in the clade (guan), and roughly eight times as rapidly as chicken. The difficulty of interpretation in this scenario was discussed by Gibbs in the context of vertebrate and poxvirus thymidine kinases [31].

### Viral IL-4 is highly conserved in ILTV

An intact vIL-4 gene was found in all 89 complete ILTV genomes published at the time of writing (after removing known recombinants, duplicates, and the original published mosaic sequence). After further removal of two entries containing unknown (N) stretches in the coding sequence, 72 vIL-4 amino acid sequences were identical to 1874C5 and Laryngo-Vac. The other 15 sequences clustered into four groups that could be distinguished by only six amino acid substitutions (Figure 6).

**Figure 6:**
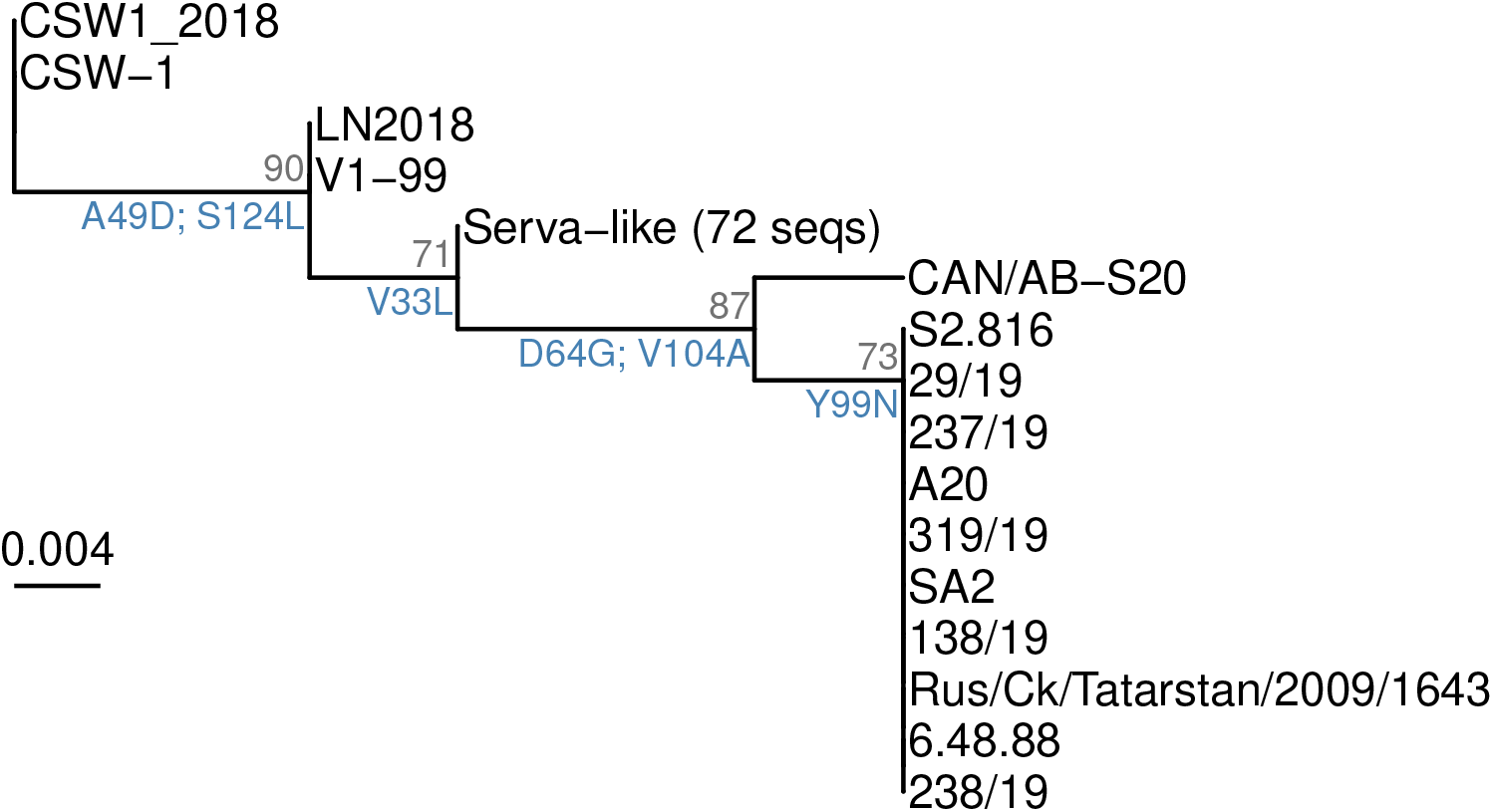
ML tree of amino acid sequences of vIL-4 in sequenced ILTV genomes. An intact gene structure and inframe coding sequence was found in all 89 complete ILTV genomes examined (after removing known recombinants, duplicates, and the original published mosaic genome). Most of the amino acid sequences were identical, and are shown as a single ‘Serva-like’ cluster. Only a handful of variants characterized the remaining sequences. Each branchpoint on the tree represents one or several amino acid variants, labeled in blue. The tree was built with IQTREE using the ‘Q.bird’ model as selected by ModelFinder, ultrafast bootstrapping (10000 iterations). Support values shown in gray are UFBoot estimations.

### Viral IL-4 as a putative IL-4/IL-13 mosaic

While vIL-4 has the highest global identity and similarity within the known protein database to avian IL-4 homologs, it also shares residue-level homology with IL-13 (Figure 7B,E). In fact, it was observed that vIL-4 has higher amino acid identity and similarity to both cIL-4 and cIL-13 than either of these cytokines have to each other (Figure 7H). In localized regions, most notably the second and fourth a-helices, vIL-4 has higher identity to cIL-13 than cIL-4 (Figure 7B).

**Figure 7:**
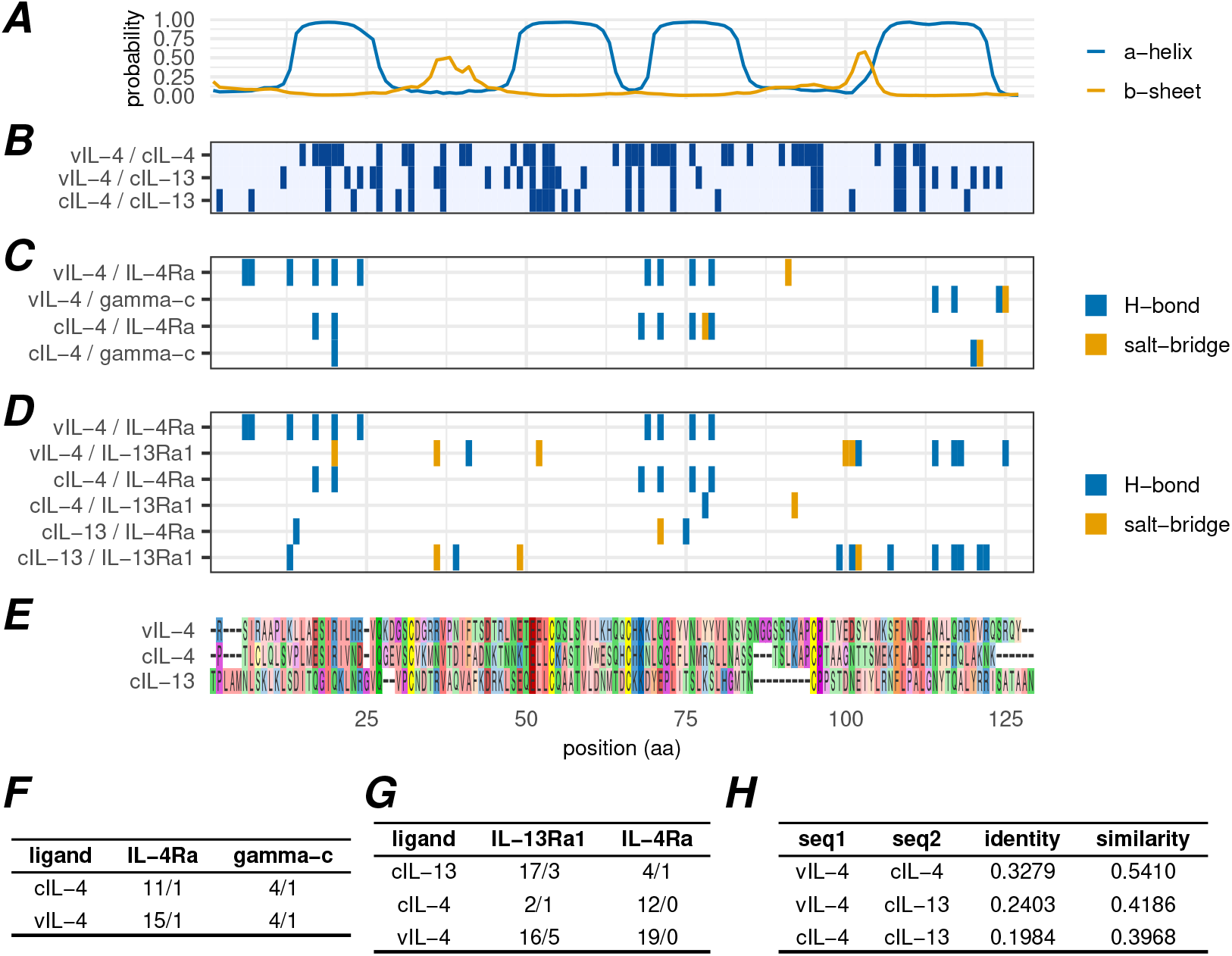
Comparison of vIL-4, cIL-4 and cIL-13 amino acid similarity and predicted receptor interactions. Panels A-E are aligned on amino acid position. (A) Secondary structure probability of the vIL-4 amino acid sequence as predicted by JPRED2. (B) Positions of identical amino acids between indicated aligned sequence pairs. (C) Positions of predicted atomic interactions between indicated ligand and IL-4 type I receptor subunit pairs (based on AlphaFold multimeric structural predictions). (D) Positions of predicted atomic interactions between indicated ligand and IL-4 type II receptor subunit pairs (based on AlphaFold multimeric structural predictions). (E) Amino acid alignment of vIL-4, cIL-4, and cIL-13 without signal peptides. (F) Number of predicted hydrogen bonds (first value) and salt bridges (second value) between two cytokine/virokine ligands and each type I receptor subunit. (G) Number of predicted hydrogen bonds (first value) and salt bridges (second value) between three cytokine/virokine ligands and each type II receptor subunit. (H) Calculated fraction identity and similarity (see methods for calculation) between each pairwise set of cytokine/virokine sequences.

IL-4 and IL-13 interact with multiple overlapping receptors, and there are two heterodimeric receptor complexes bound by IL-4: the type I complex comprises IL-4R α and common gamma chain (γC) subunits while the type II complex comprises IL-4Rα and IL-13R α1 subunits [32]. IL-13 also binds the type II complex, in addition to its own separate IL-13Rα2 monomeric receptor. IL-4 and IL-13 interact with the type II receptor with different modes and affinities: while high-affinity binding of IL-4 to IL-4Rα is followed by recruitment of IL-13Rα1, IL-13 binds most strongly to IL-13Rα1 with subsequent IL-4Rα recruitment [32]. Given the observed localized regions of higher identity between vIL-4/cIL-4 and vIL-4/cIL-13, multimeric structural predictions were generated using AlphaFold to investigate the potential impact of these localized differences on predicted protein-protein interactions.

The positions of predicted hydrogen bonds and salt bridges between each combination of potential ligand and receptor subunit were calculated for both type I and II receptors (Figures 7C and D). Both cIL-4 and vIL-4 are predicted to interface with the type I receptor subunit IL-4Rα along the first and third α-helices and with γC within the fourth helix. Of particular note, however, are the predicted interactions with the type II receptor. In line with known binding affinities described above, cIL-4 is predicted to interact primarily with IL-4Rα, while cIL-13 is predicted to interact primarily with IL-13Rα1. In contrast, vIL-4 is predicted to form a large number of hydrogen bonds with *both* IL-4Rα and IL-13Rα1, with the interacting regions positionally homologous to those of the cellular cytokine/subunit interactions. These interactions are quantified in 7G. It is important to note that all of the multimeric AlphaFold predictions exhibited low predicted aligned error (PAE) scores between all chains, indicating confident relative positioning, with the exception of the IL-4/IL-13Rα1 interface (Figure S4). Despite repeated modeling attempts, this interface was never modeled with high confidence; it is therefore possible that the predicted interactions for this pair are underestimated.

### Functional equivalence of vIL-4

We investigated the ability of both cIL-4/GFP and vIL-4/GFP to stimulate the production of nitric oxide (NO) in avian macrophage cells. Macrophages play a critical role in the immune system and are known to produce NO in response to certain stimuli. For this experiment, the avian macrophage cell line HD11 was utilized. Cells were stimulated with varying concentrations (ranging from 62.5 to 1000 ng/ml) of the tagged vIL-4 and cIL-4 proteins for a duration of 4 hours. This stimulation period allowed for interaction between the tagged proteins and their cognate receptors on the macrophage cell surface and the triggering of downstream signaling pathways. Subsequently, *E. coli* lipopolysaccharide (LPS), a potent bacterial endotoxin known to stimulate NO production in macrophages, was added to the cultures to activate the macrophages. Following a 20-hour incubation period, the concentration of nitrite (NO_2_−) in the cell culture supernatant was measured using the Griess assay. Both chicken interleukin-4 (cIL-4) and viral interleukin-4 (vIL-4) exhibited similar stimulatory effects on nitric oxide (NO) synthesis in macrophages when co-stimulated with the microbial agonist lipopolysaccharide (LPS) (Figure 8). No statistically significant difference was observed in the amount of nitrite (NO2-), a stable breakdown product of NO, produced by macrophages treated at each concentration of either cIL-4 or vIL-4 in the presence of 2.5 μg/mL LPS. This suggests a functional equivalency between the two interleukin-4 analogues in their ability to induce NO production in macrophages upon LPS stimulation [33–36].

**Figure 8:**
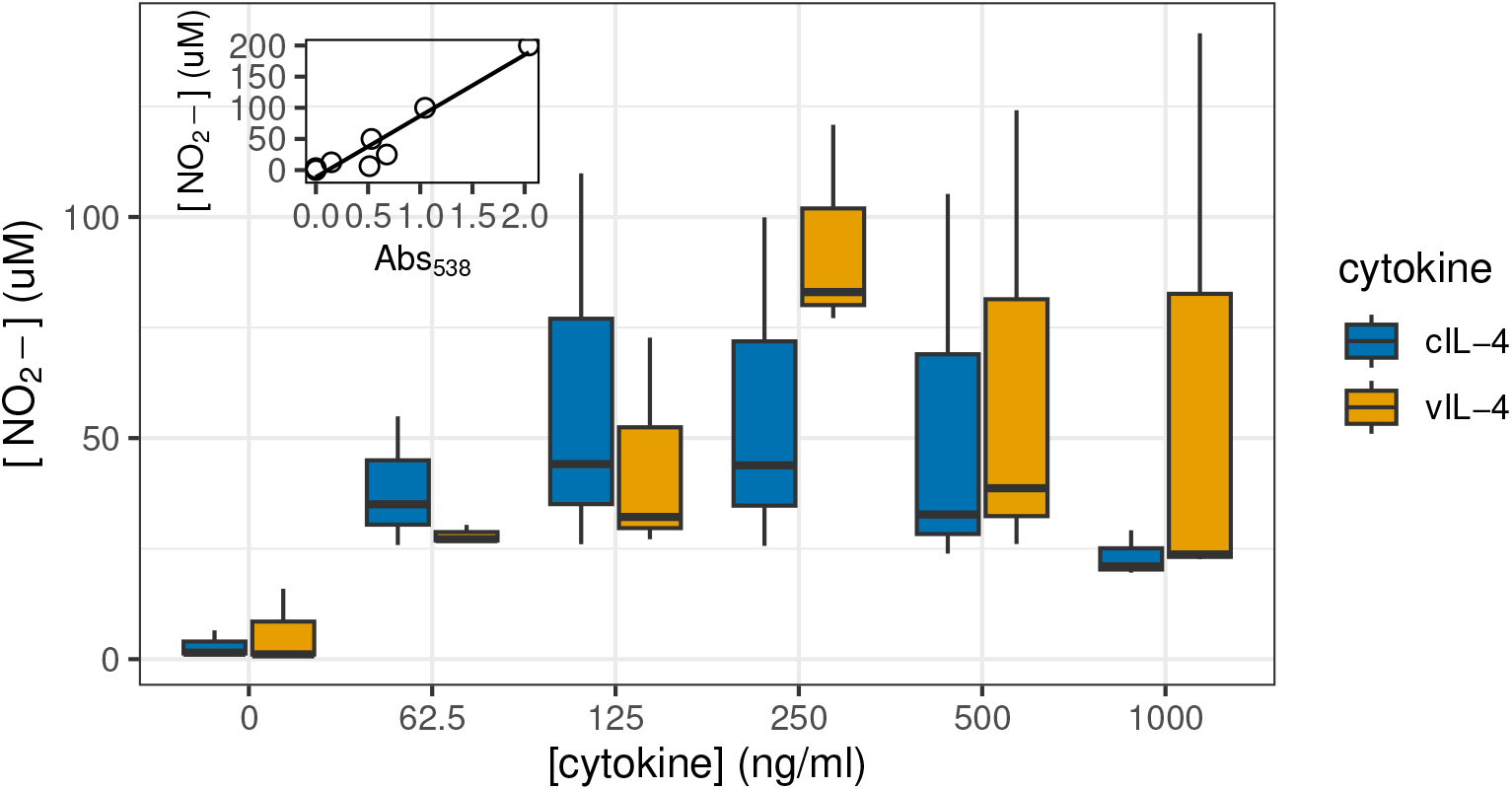
Stimulation by IL-4 of nitric oxide production in macrophages. The Griess assay was used to measure NO_2_− as an indicator of NO production at varying concentrations of either chicken IL-4 (cIL-4) or viral IL-4 (vIL-4) and in the presence of 2.5 μg/ml LPS. Inset: standard curve of Griess reagent used to calculate NO_2_− concentrations (R^2^= 0.92).

### In vitro *replication kinetics of parent 1874C5 and ΔvIL4 strains*

The replication kinetics of parent 1874C5 and ΔvIL-4 ILTV strains were determined by measurement of TCID_50_ /ml (Figure 9A) and viral genome load (Figure 9B) from combined infected supernatant and cells collected at 24, 48, 72 and 96 hours post virus absorption. No measurable cytopathic effect was detected at 24 hours post-absorption for either 1874C5 or ΔvIL-4 strains. By 48 hours post-absorption, the parent 1874C5 strain had an average titer of log_10_ 0.24 TCID_50_/ml as compared to an average titer of log_10_ 2.99 TCID_50_/ml for the ΔvIL-4 strain. At 72 and 96 hours the ΔvIL-4 strain titers were log_10_ 1.1 to 1.7 TCID_50_/ml higher than the parent 1874C5 strain that reached an average titer of log_10_ 4.40 TCID_50_/ml by 96 hours post-absorption. Despite the difference in viral TCID_50_/ml, viral genome load for both viruses showed an almost equivalent rate of increase by 72 and 96 hours post-absorption (Figure 9B).

**Figure 9:**
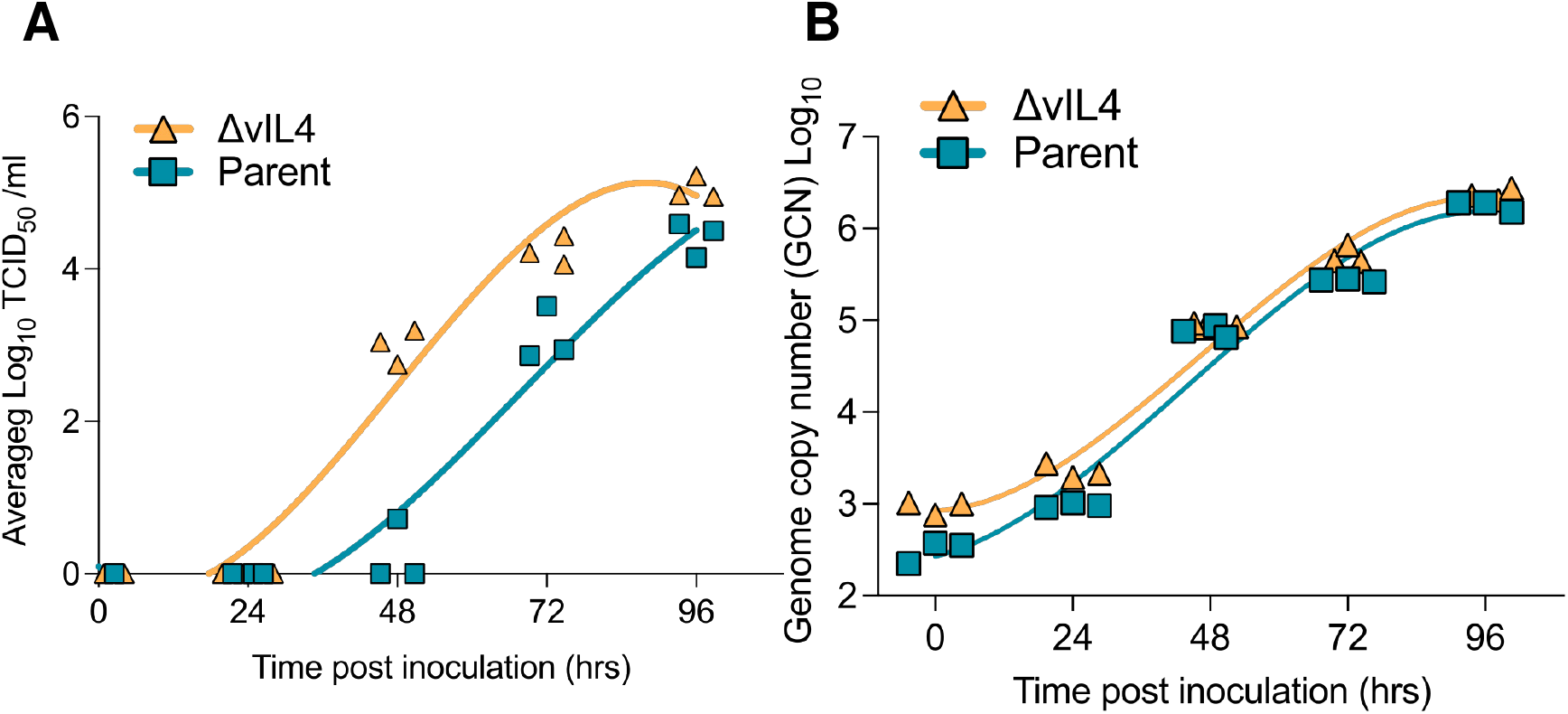
Growth kinetics of 1874C5 parental and ΔvIL-4 ILTV strains in the LMH cell line. LMH cells (1.4×10^6^ cells per well) were inoculated in triplicate with 1874C5 parental strain and ΔvIL-4 strain at MOI of 0.002. After absorption for 1 hour at 39°C and 5% CO_2_, cells were washed and incubated for 0, 24, 48, 72 and 96 hours. At each time point supernatant and cells were collected (n=3) and (A) viral titer and (B) viral genome copy number were determined for each strain at each time point. Individual replicates and corresponding nonlinear fit curves are shown.

### Pathogenicity of parent 1874C5 and ΔvIL4 strains after intratracheal inoculation

The percent survivability of chickens inoculated with the ΔvIL-4 strain (n=17) was higher (47.1%) than the survivability of chickens (n=15) inoculated with the parent 1874C5 strain (33.3%). Mortalities induced by the parent 1874C5 strain peaked at day four, while mortalities for chickens inoculated with the ΔvIL-4 strain peaked at five d.p.i. (Figure 10A). Average clinical sign scores (CSS) induced by the parent 1874C5 and ΔvIL-4 strains at days 2 to 7 p.i. were not statistically different (Figure 10B). However, at days 4 and 5 p.i. clinical sign scores induced by the parent 1874C5 strain (4.60 and 5.00) were numerically higher than clinical signs scores induced by the ΔvIL-4 strain (3.55 & 3.89). The peak of clinical signs for both the parent 1874C5 and ΔvIL-4 groups of chickens were identified at day five p.i. (Figure 10B) which coincided with the ΔvIL-4 strain peak of mortality. However, the peak of mortality for the parent 1874C5 inoculated group of chickens was recorded earlier (day 4 p.i.) than for the ΔvIL-4 strain peak of mortality (day 5 p.i.) (Figure 10A). The average tracheal viral genome load for the parent 1874C5 and ΔvIL-4 strains was measured at 3 and 5 d.p.i — no statistical differences were detected. (Figures 10C & D). At three days pi the parent 1874C5 and the ΔvIL-4 viral genome load in trachea reached an average of 5.339 and 5.773, respectively (Figure 10C). At 5 days p.i., although not statistically different (p>0.05), the average viral genome load for parent 1874C5 strain was higher (4.6) than that of the ΔvIL-4 strain (3.3) (Figure 10D). When comparing decreases of viral genome load of each strain in the trachea between days 3 and 5 p.i., the parent 1874C5 genome load showed no significant decrease (p>0.05) (Figure 10E) while the genome load of ΔvIL-4 strain showed a significant decrease (p=0.001) from 5.8 at day 3 to 3.3 at day 5 p.i. (Figure 10F).

**Figure 10:**
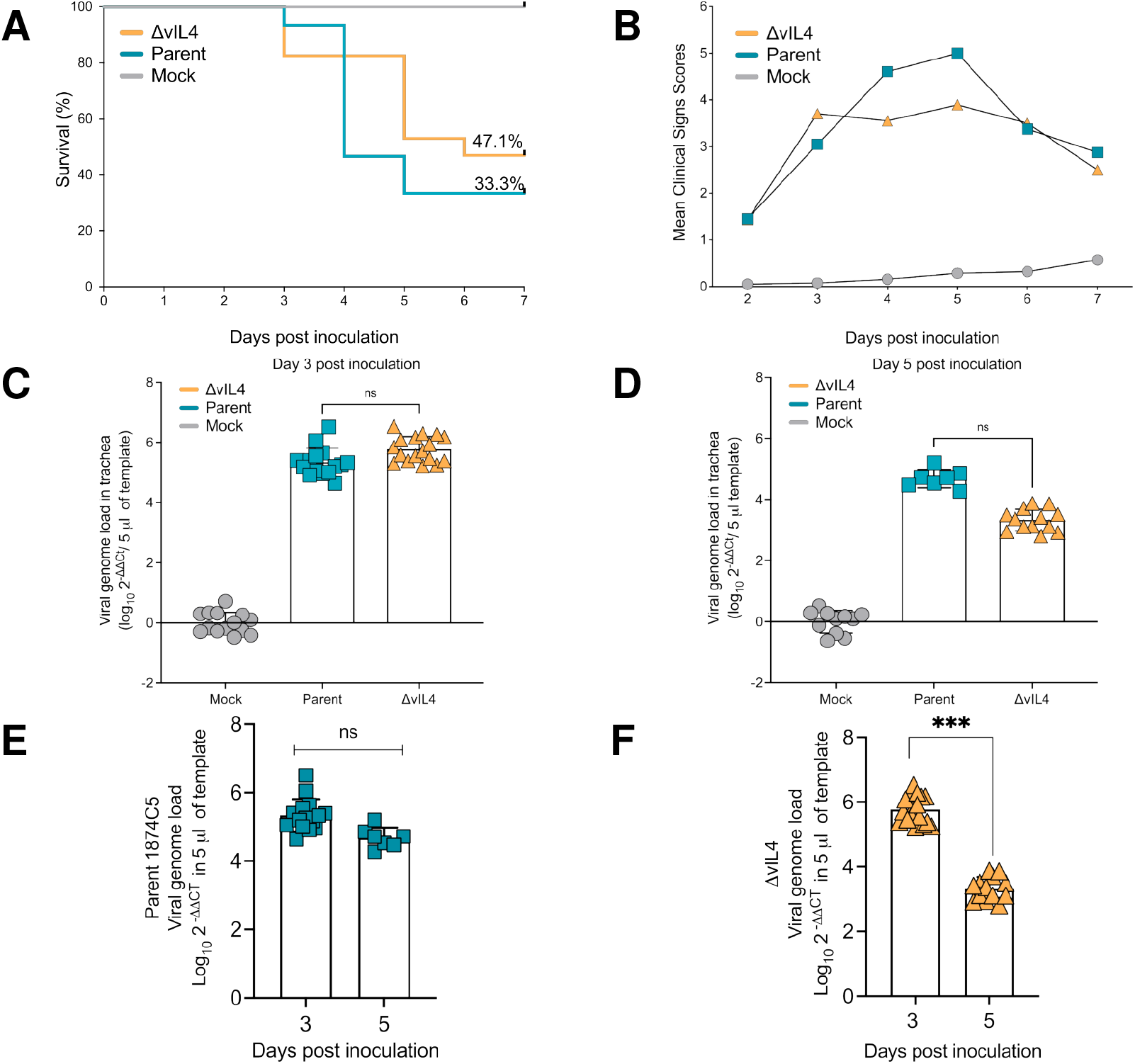
Pathogenicity of 1874C5 (parent) and vIL-4 gene-deleted strain (ΔvIL-4) in specific pathogen free (SPF) chickens after intratracheal inoculation at 24 days of age. (A) Percent survival for chickens inoculated with the ΔvIL-4 strain (n=17) was 47.1% and for chickens inoculated with the parent strain (n=15) was 33.3%. (B) Average clinical sign scores (CSS) per group of chickens from 2 to 7 days post-intratracheal inoculation. Although not significant, average CSS induced by the 1874C5 strain were higher than average scores induced by the knockout. (C) Viral genome load in trachea three days post-intratracheal inoculation. (D) Viral genome load in trachea five days postintratracheal inoculation. (E) Comparison of parent 1874C5 genome load at 3- and 5-days post-inoculation. (F) Comparison of ΔvIL-4 genome load at 3- and 5-days post-inoculation. Individual replicate loads indicated by points; column heights indicate the average trachea viral genome load per group. No significant (ns) differences (p>0.05) were detected between viral genome load of the 1874C5 parent and the ΔvIL-4 mutant at either 3- or 5-days post-inoculation. Significant differences (p<0.05) were detected for the average viral genome load of ΔvIL-4 in the trachea at 3-vs 5-days post-inoculation.

Based on the Guy et al 1990 histopathological trachea lesion score method [37], all upper tracheas from the parental-inoculated group of chickens received the highest score of five while all trachea sections from the ΔvIL-4-inoculated group of chickens received a score of four. Trachea ring sections from the parental-inoculated chickens displayed detachment of large segments of the mucosal epithelium and the submucosa with syncytia cells and inclusion bodies rarely found and fibrino-haemorrhagic cellular infiltrates released into the trachea lumen. The score five tracheas are characterized by the absence of any epithelial mucosal cells, with areas where lean layer of submucosa remains (Figure 11A), and areas where enlarged submucosa is still present (Figure 11B). Although the ΔvIL-4-inoculated upper trachea sections displayed damage and detachment of the mucosal epithelium, trachea segments maintained a large portion of the submucosa and in some instance distal thin mucosal epithelium with scattered syncytia and intranuclear inclusion bodies could still be detected (Figure 11C; see inserted photomicrograph). Another histopathological presentation frequently found in the tracheas of the ΔvIL-4-inoculated chickens was the presence of slightly hemorrhagic cellular pseudo-membranes that remained associated to the enlarged submucosa (Figure 11D).

**Figure 11:**
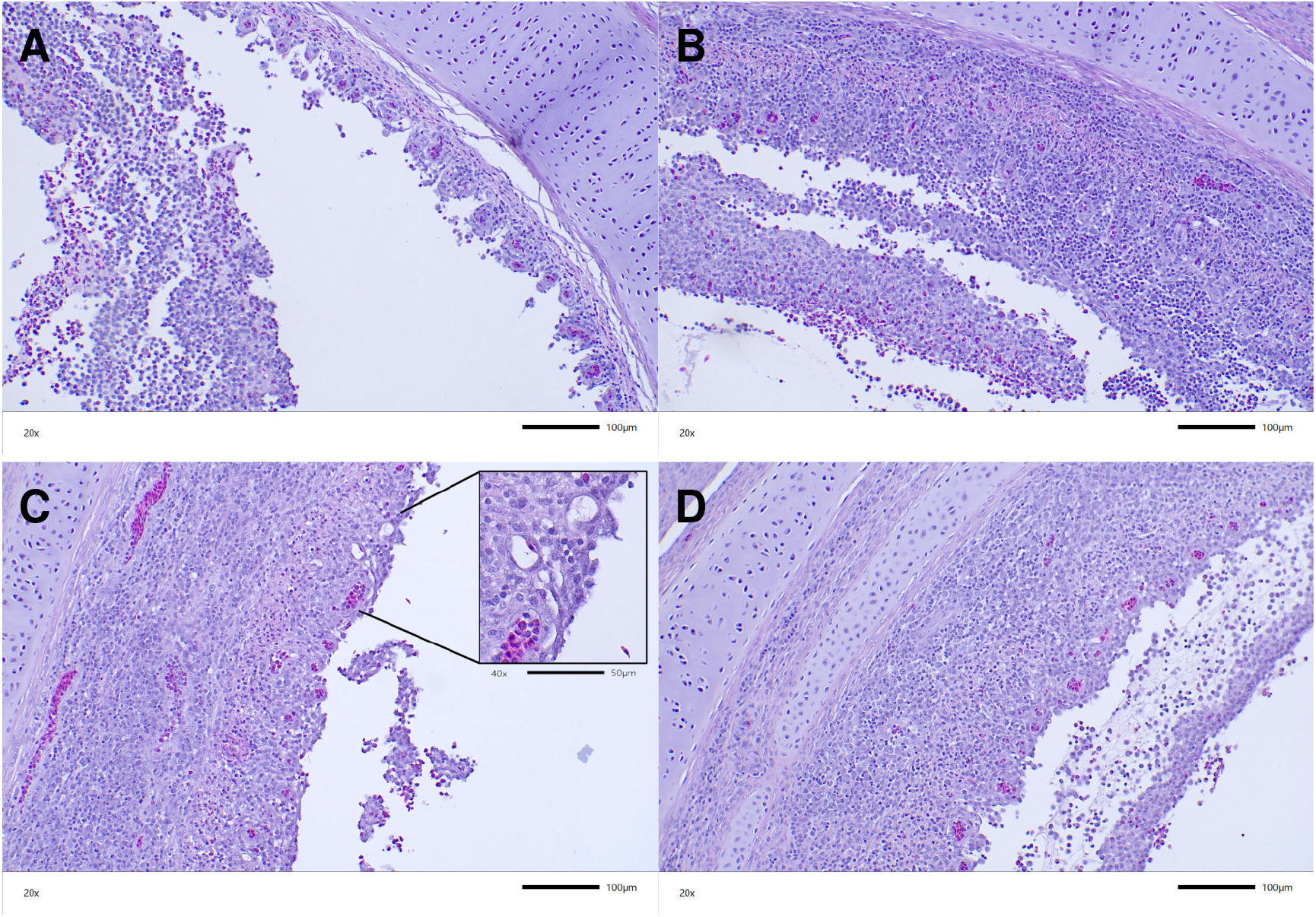
Histopathological lesions in the trachea induced by parental 1874C5 and ΔvIL-4 at four days post-intratracheal inoculation. Photomicrographs are H&E-stained upper trachea sections from chickens infected with parental 1874C5 (A & B) or ΔvIL-4 (C & D) at 20x magnification with image scale bar of 100 μm. (C) inset: Inserted photomicrograph at 40x magnification with image scale bar of 50 μm; underscores the presence of mucosal epithelium and goblet cells with few syncytia and intranuclear inclusion bodies still attached distal of the submucosa.

## Discussion

Vertebrate immune systems have evolved intricate mechanisms to skillfully recognize and control viral infections. However, viruses have developed countermeasures to subvert host immunity and ensure their continued survival. A key driver in this host-pathogen co-evolution is horizontal gene transfer, in which large DNA viruses acquire host genes. Over time, these captured genes are modified through mutation and selection to perform four main functions: (i) mimicry of host molecules to evade detection; (ii) antagonism of immune pathways for direct suppression; (iii) development of novel immunoregulatory function to manipulate the host response; or (iv) stimulation of immune target cells to facilitate virus replication or latency. Notably, the acquisition and subsequent adaptation of antiviral defense components confer selective advantages on the virus to directly target and disrupt both the innate and adaptive arms of the immune response, ultimately enhancing virus replication and dissemination. Interleukins play a fundamental role in regulating the magnitude of protective responses. Accordingly, several herpesviruses encode functional homologs that “mimic” host cytokines (IL-6, IL-10, and IL-17). Virokines are more common in γthan β-herpesviruses [38] with no reports of non-chemotactic cytokines encoded by α-herpesviruses prior to this discovery of an interleukin 4 mimic in the genome of ILTV. One chemokine, vIL8, is encoded by the related avian α-herpesvirus Gallid alphaherpesvirus 2 (GaHV-2), the causative agent of Marek’s disease [39]. Generally, virokines can interact with immune cell receptors and activate pathways that play a role in the pathogenesis of herpesvirus-associated diseases by modulating immune responses, promoting cell survival, and stimulating cell proliferation.

One of the most well-studied virokines is viral interleukin-6 (vIL-6), encoded by Kaposi’s sarcoma-associated herpesvirus (KSHV). Despite sharing only 37% amino acid identity with human IL-6, vIL-6 exhibits potent pro-inflammatory activity. It acts in an autocrine manner, promoting both cell growth and survival by inducing expression of host IL-6 [40] and suppressing production of IFNa, a key molecule for activating NK cells and macrophages and enhancing their anti-viral activity.

A striking convergence in viral evolution is evident in the independent acquisition of interleukin 10 homologs by more than a dozen viruses belonging to three distinct viral families: eight in Herpesviridae, one in Alloherpesviridae and four in Poxviridae [41]. The widespread distribution of interleukin-10 (IL-10) mimics among diverse viral families highlights a profound selective pressure during viral-host co-evolution to acquire genes or cDNA copies encoding proteins capable of immunosuppression. The best-studied IL-10 homologs are those of EpsteinBarr virus (EBV) and cytomegalovirus (CMV) [42]. The gene encoding vIL-10 of EBV has over 80% amino acid identity with the human counterpart. Like vIL-6 of KSHV, EBV’s vIL-10 inhibits IFN-γ synthesis in T cells [43] and potentially inhibits natural killer (NK) cells and cytotoxic T cells, as evidenced by reduced activities of these cells in mice infected with a recombinant vaccinia virus expressing vIL-10 of EBV [44]. This suppression hinders the immune system’s ability to clear the virus and potentially promotes viral persistence within the host. In addition, vIL-10 retains its ability to stimulate B cell growth to facilitate virus replication in these cells [45,46].

Cytomegaloviruses of primates have also adapted to manipulate the host immune response through an IL-10 homolog [47–49]. The protein cmvIL-10 in HCMV arose from a captured human IL-10 gene some >42 million years ago, evolving a molecule with only 27% amino acid sequence identity to its human counterpart. Despite this low homology, cmvIL-10 retained the ability to bind the human IL-10 receptor with high affinity [50] and exerts a multifaceted immunosuppressive effect depending on the target cell and the stage of infection.

Saimiriine gammaherpesvirus 2 (SaHV-2) further exemplifies viral mimicry with its unique IL-17 mimic, vIL-17 [12,51]. This virokine exhibits a high degree of structural similarity (74% amino acid identity) to its host counterpart and can bind the same cellular receptor [12]. This receptor is present on T cells, B cells, fibroblasts and other cells types, and vIL-17 binding to the receptor may stimulate metabolism, differentiation or proliferation to facilitate virus replication [12] or manipulate the immune response to favor virus escape.

These observations collectively reinforce the broader concept that virally encoded cytokine mimics represent a strategic advantage for herpesviruses, depending on the mimicked function, to replicate and establish persistent infections within their hosts. Anti-inflammatory mimics like vIL-10 directly suppress the host immune response, promoting viral persistence. Even proinflammatory mimics can provide benefit to the virus by stimulating growth of potential virus target cells or by inhibiting diverting responses from antiviral immune pathways. Viral IL-6, while triggering inflammation, suppresses antiviral interferon production and promotes cell survival, potentially aiding the virus. Similarly, saimiriine herpesvirus 2’s vIL-17 might enhance virus replication in target cells or alter antiviral immunity. Overall, the benefit hinges on how the mimic manipulates the immune response.

Here, we report the discovery of a functional interleukin-4 (IL-4) homolog in an α-herpesvirus, adding to the list of known virokines and uncovering a previously undocumented immunosuppressive tactic employed specifically by an avian herpesvirus. In mammals, interleukin-4 (IL-4) acts as a central promoter of type-2 immunity. It influences a variety of cells, both haematopoietic and non-haematopoietic. Its effects include activating specific Th2 cells in an autocrine fashion; differentiating B cells into plasma cells; promoting immunoglobulin E (IgE) class switching; generating alternatively activated macrophages; stimulating smooth muscle contraction, mast cell activation, and mucus production; innate cell recruitment to the site of inflammation; blocking the generation of Th1- and IFN-γ-driven responses; and inhibiting cell-mediated immunity by diverting the function of CD8 T cells away from antiviral cytolysis [52]. Research on the functional role of chicken IL-4 is less comprehensive compared to its mammalian counterpart. Avian species, in contrast to mammals, lack the immunoglobulin E (IgE) class, but IL-4 likely promotes expression of IgY and IgA antibodies. Notably, elevated levels of IL-4 expression have been observed in vaccinated chickens [53,54].

A transcript likely representing vIL-4 was previously identified by Northern blot with a probe covering the UL/IR_s_ junction forward strand [55]. The 0.7 kb transcript referred to as “IR1” was mentioned in passing as being non-polyadenylated with late kinetics, and was suggested as a candidate for further study (the primary vIL-4 transcript described here is 634 nt and in roughly the same location and orientation, although it was polyadenylated). Due to the extensive splicing of this transcript and the limitations of early protein databases and search tools, it appears that no further research was conducted on this transcript until the present study. Computational methods at the time likely failed to identify potential protein-coding reading frames within the region, hindering its recognition as a functional transcript.

*In vivo* RNA-Seq showed vIL-4 to be among the top five most abundantly transcribed viral genes during the sampling period from 3 to 7 days post-infection, which encompassed the period of virulent lytic replication. Translation of the vIL-4 protein was subsequently confirmed using tandem mass spectrometry of infected cell culture and was absent in uninfected cells. In cells infected with both virulent and attenuated strains of ILTV, vIL-4 protein expression ranked in the top third of detectable viral proteins by relative molar abundance, and 86% of the predicted detectable mature protein sequence was covered by experimentally detected peptides. From this evidence, and from a retrospective look at previously published expression data as described herein, it is reasonable to conclude that transcription and translation of vIL-4 is ubiquitous, though variable, across a range of cell types and stages of lytic infection. Additional targeted and controlled studies will be necessary to further characterize the kinetics of vIL-4 expression and the limits of tissue specificity, if any.

Our analysis demonstrates that the four-exon genomic structure of vIL-4 is remarkably conserved between chicken and other vertebrate IL-4 homologs and the amino acid sequence displays structural conservation with vertebrate homologs based on computer modeling. Phylogenetic analysis suggests its direct capture from a Galliformes host approximately 75 Mya. The Iltovirus genus has been previously estimated to have diverged from between 200 and 90 Mya [56,57], and the last common ancestor between ILTV and Psittacid alphaherpesvirus 1 (PsHV-1), the most closely-related sequenced Iltovirus, was recently estimated at between 30 and 50 Mya [57]. PsHV-1 has a similar gene arrangement at the UL/IRS junction, and also contains a ~2.4 kb unannotated region between UL(−1) and ICP4. However, no gene structure or coding region homologous to vIL-4 was found in this region in PsHV-1 based on searches using BLASTN, TBLASTN, protein-to-genome alignment with `miniprot`, and *de novo* gene prediction with AUGUSTUS. The only other published genome from a virus assigned to the *Iltovirus* genus, PsHV-5 [58], was similarly searched and no vIL-4 homolog was identified. It is notable that another avian herpesvirus from a different genus, Gallid alphaherpesvirus 2, encodes a protein (vIL-8) with homology to the vertebrate C-X-C chemokine interleukin 8 [39]. Like vIL-4 in ILTV, this viral gene has conserved exonic structure with the homologous host gene. It is located near the terminal repeat junctions of the genome and thus, like vIL-4, near the end of the fully processed viral genome, suggesting it may have been captured from the host by a similar mechanism.

Functionally, vIL-4 stimulates nitric oxide production in macrophages to levels comparable to chicken IL-4. Notably, compared to the parent 1874C5 strain, an ILTV mutant lacking vIL-4 (ΔvIL-4) exhibited slightly higher viral titers but similar genome load in LMH cells. This result may suggest that *in vitro*, the absence of vIL-4 may favor a faster production of infectious viral particles, reflected by the increased early titers for the ΔvIL-4 strain. In contrast, the rate of viral genome replication *in vitro* was similar for both strains, which coincides with the *in vivo* results in which parent 1874C5 and ΔvIL-4 viral genome load in the trachea at 3- and 5-days p.i. showed no difference. However, when comparing the trachea infected with either viral strain, the genome load of ΔvIL-4 was significantly lower at day 5 pi. Also, the ΔvIL-4 strain induced lower mortalities and lower clinical signs of disease. These subtle differences in pathogenicity were corroborated by the histopathological findings in the upper trachea of chickens inoculated with the ΔvIL-4 strain. While the tracheal lesions induced by the ΔvIL-4 strain were characterized by increased infiltration and epithelial damage, in most of the trachea rings examined the submucosa and lamina propria remained in place with some distal epithelium still attached. In the case of 1874C5-inoculated chickens, most of the trachea mucosa and submucosa were detached into the lumen. These findings collectively suggest that vIL-4 acts as a novel virokine, potentially modulating the host immune response and contributing to ILTV virulence.

The discovery of vIL-4 with sequence similarity to cIL-4 also raises intriguing questions about the connection between this viral mimic and other cytokines, particularly cIL-13. In mammals, both IL-4 and IL-13 activate common pathways through usage of lineage-determining transcription factors STAT6 and GATA3 following binding to shared receptor subunits. Viral IL-4 might, therefore, not only mimic cIL-4 but potentially emulate some functions of chicken IL-13 as well. Supporting this hypothesis, vIL-4 shares localized sequence similarity to *both* cIL-4 and cIL-13, and structural prediction suggests that vIL-4 may bind both type I and type II IL-4 receptor dimers with higher affinity than either cellular cytokine. Empirical binding studies are needed to validate these predictions. However, collectively these findings suggest vIL-4 and cellular IL-4/IL-13 may act in concert to manipulate the host immune response, diverting it away from a protective Th1 response that favors cytolytic CD8 T cells and antiviral immunity and toward a Th2 response. Although this Th2 response, through the production of matrix metalloproteinases and goblet cell metaplasia leading to mucus hypersecretion and mast cell activation, may be beneficial to the virus in the short term, the extensive mucus production can ultimately lead to asphyxiation, the primary cause of death in ILTV infection.

In conclusion, this study presents the discovery of a previously unknown vIL-4 encoded by ILTV. It is likely that ILTV has developed a multifaceted approach to manipulate the host’s immune response through the action of its encoded cytokines, glycoprotein G (chemokine) and vIL-4 (virokine). The vIL-4 virokine may play a role in ILTV virulence, potentially through interactions with both IL-4 and IL-13 receptors. Paradoxically, vIL-4 might even promote specific aspects of the humoral immune response that favor the virus through a shift away from the protective Th1 responses toward Th2 responses. Further investigation into the complex interplay between vIL-4, cIL-4, cIL-13 and their receptors should pave the way to a better understanding of immune evasion strategies employed by ILTV, undoubtedly aiding efforts to develop more effective strategies to combat infectious laryngotracheitis.

## Data availability

Sequences for the 1874C5 and Laryngo-Vac vIL-4 mRNA transcripts have been deposited in GenBank under accession numbers PP974316 and PP974317. Raw short- and long-read sequencing data is available in the NCBI SRA under BioProject PRJNA1167273. Raw LC-MS/MS data has been deposited in the public MassIVE repository (https://massive.ucsd.edu) under project accession MSV000097080 (ProteomeXchange dataset PXD060595). All curated nucleotide and amino acid sequences referenced, along with GFF3 annotations for homologs in public vertebrate and ILTV genomes, are available at the manuscript GitLab repository (https://gitlab.com/BASE2BIO/projects/vIL4). The public git repository also includes all code used to perform statistical analyses, generate figures, and assemble the manuscript in RMarkdown.

## Materials and Methods

### Viruses and Cells

The ILTV virulent strain 1874C5 was originally isolated from the upper respiratory tract of broilers during outbreaks of the disease and subsequently classified as a member of the genotype VI [59]. The CEO vaccine strain Laryngo-Vac was a generous gift from Zoetis. The vIL-4 null mutant was derived from the genotype VI virulent 1874C5 strain. Leghorn male hepatoma (LMH) cells were propagated in Dulbecco’s Modified Eagle Medium (DMEM) supplemented with 10% fetal bovine serum (ThermoFisher), 100 U/ml penicillin, 100 μg/ml streptomycin and 2mM L-glutamine. For virus infections, maintenance DMEM supplemented with 2% FBS was used for the generation of vIL-4 null mutant and replication kinetics studies. ILTV strains CEO, 1874C5 and ΔvIL-4 virus were also propagated in primary chicken kidney (CK) cells from 3-to 4-week-old chickens for the purpose of generating virus stocks and isolation of nucleic acids as previously described [60,61]. Titers were expressed as tissue culture infective dose (TCID_50_) and calculated by the Reed and Muench method [62].

Avian macrophage cells (HD11) were cultured in DMEM supplemented with 10% fetal bovine serum, 100 U/ml penicillin, 100 μg/ml streptomycin, and 2.0 mM L-glutamine. For LPS/IL-4 stimulation assays, aliquots of 100 microliters of cell suspension, containing 2×10^6^ cells/ml, were seeded into wells of a 96-well tissue culture plate. The cells were propagated until they reached approximately 85% confluence before further experimentation.

### RNA isolation

Chicken kidney (CK) cells were infected with 1874C5 or LaryngoVac at an MOI of 0.001. RNA was isolated from infected CK cells using Trizol according to the manufacturer’s recommendations. RNA was quantified using the Qubit RNA BR (Broad-Range) Assay Kit on a Qubit Fluorometer (ThermoFisher). The quality of RNA was assessed using the Agilent 2100 Bioanalyzer (Agilent Technologies) with the RNA 6000 Nano LabChip kit according to the manufacturer’s instructions. Briefly, the RNA samples were diluted in water and loaded onto the microfluidic chip along with RNA ladder standards. The resulting electropherograms were analyzed with Agilent 2100 Expert software to determine the RNA Integrity Number (RIN).

### PacBio sequencing

RNA samples with RIN ≥ 7 were used to prepare the IsoSeq barcoded libraries following the PacBio protocol (Procedure & Checklist – Iso-Seq Express Template Preparation for Sequel and Sequel II Systems, PN 101-763-800 Version 02) using the SMRTbell Express Template Prep kit 2.0 (PacBio). The individual libraries were barcoded using the Iso-Seq Express Oligo kit (Pacbio PN 101-737-500). The barcoded libraries were multiplexed in equimolar amounts into a final SMRTbell template mix, which was purified using 1×beads. The final library was sequenced on the Sequel II system for 20 hours using the Sequel II Binding Kit 2.1. The final loading concentration was 80 pM.

### IsoSeq preprocessing and transcript building

Raw PacBio raw subreads were processed to circular consensus sequencing (CCS) reads using ccs v6.4.0 (`--min-rq 0.9`) (this and all other PacBio software referenced was installed from the PacBio Bioconda packages (https://github.com/PacificBiosciences/pbbioconda) [63]. Demultiplexing and adapter removal was performed using lima v2.7.1 (`--isoseq–peekguess`). Poly-A tail removal and concatemer filtering were performed using isoseq refine from the isoseq3 package v3.8.2 (`--require-polya`). Full-length, non-concatemeric (FLNC) reads were then clustered into non-redundant transcripts using isoseq cluster (`--use-qvs`). Transcript cluster sequences were mapped against the combined bGalGal1.mat.broiler.GRCg7b+virus genome reference using pbmm2 align v1.10.0 (`--preset ISOSEQ–sort`). Overlapping, exon-sharing transcripts were further collapsed using isoseq3 collapse (`--do-not-collapse-extra-5exons– max-5p-diff 200`). Only transcripts with a minimum support of four FLNC reads were kept. Further filtering of potential artifacts was performed using SQANTI3 v5.1.1 [64] to remove transcripts with a genomic adenine content downstream of the mapped TTS of 60% or greater (suggesting artifactual priming on the genome rather than a transcript poly-A tail) and spliced transcripts with non-canonical donor/acceptor sites. IGV [65] was used to visualize transcripts on the viral genome to first detect the presence of the vIL-4 transcript cluster.

### Curation of IL-4 homologs

Initial BLASTP and TBLASTN [66] searches of the chicken IL-4 sequence (NP_001385388) against the *nr* and *nt* databases, respectively, were used to generate a candidate list of published homologs. In some cases, high-confidence annotations were found for a species; in others, only partial TBLASTN matches to genomic regions were seen. In the latter case, the chromosome/scaffold/contig sequence was analysed using a combination of miniprot [67] protein-to-nucleotide alignment with GFF3 output and AUGUSTUS [68] *de novo* gene prediction (*Gallus gallus* model) to generate a preliminary gene structure. After multiple alignment of these preliminary sequences, several homologs from existing database entries contained questionable insertions/deletions in conserved regions. These homologs were analyzed either visually in IGV or using miniprot alignment to see if valid adjustments could be make to the splice junctions. In all such cases, valid adjustments using canonical donor/acceptor sites were found that brought the protein sequences into better structural agreement with the group as a whole. The final gene models in GFF3 format are available in the manuscript repository as described in *Data Availability*. Notes on the curation of each homolog sequence can be found in Table S1. Homolog sequences that were either absent from existing genome annotations or were modified from the existing annotation were submitted to GenBank as third-party annotations (TPA:inferential) with accession numbers XXX–XXX.

### Structural modeling

The JPred4 web server was used for predicting secondary structure of the vIL-4 amino acid sequence [69]. The SignalP 6.0 web server was used to predict signal peptide regions [70]. AlphaFold 2.0 [71] via ColabFold [72] (`--amber --use-gpu-relax`, all others defaults) was used for tertiary structure prediction of both vIL-4 and chicken IL-4, individually and in conjunction with the sequences of the type I and type II receptor complex (IL-4Rα + γc and IL-4Rα + IL-13Rα1, respectively). All input files used in prediction, output models and QC plots are available in the manuscript repository. ColabFold was run in the cloud using Nextflow [73] and the LocalColabFold installer (https://github.com/YoshitakaMo/localcolabfold). The PDBsum1 tool (version unknown, downloaded 2024-07-25) was used to predict hydrogen bonding and salt bridge interactions from AlphaFold structural models [74].

### Phylogenetics and tree reconstruction

MAFFT v7.525 [75] (`--auto`) was used to build all CDS and amino acid multiple alignments used in the paper. Tree inference was performed using IQ-TREE v2.3.4 [76]. Models for each multiple alignment were initially chosen using the ModelFinder feature of IQ-TREE [77] (‘Q.bird+I+G4’ for IL-4 amino acid alignment; ‘GTR+I+R’ for IL-4 coding sequence alignment; ‘Q.bird’ for ILTV vIL-4 alignment. Bootstrap support was computed using UFBoot2 with 10000 iterations [78].

### Generation of GFP/IL-4 fusion proteins

To evaluate the functional similarity between viral interleukin 4 (vIL-4) and chicken interleukin 4 (cIL-4), both proteins were N-terminally tagged with a green fluorescent protein (muGFP) [79] (Figure S2A). The amino terminus was chosen based on *in silico* spatial modeling of the IL-4 receptor complex (see Figure 4) in order to minimize potential interference with receptor binding. A flexible linker peptide consisting of 12 amino acids (GSAGSAAGS-GEF) [80] was inserted between the muGFP and the respective IL-4 coding sequences. This linker serves a crucial role in maintaining proper protein folding and minimizing steric hindrance, ensuring the tagged protein retains functionality. The vIL-4 signal peptide sequence was incorporated at the N-terminus of the muGFP moiety in both constructs. This sequence directs the tagged proteins towards the secretory pathway, enabling their efficient export from the cell. To monitor transfection efficiency, an internal ribosome entry site (IRES) was placed upstream of a separate reference fluorescent protein (red fluorescent protein; RFP). This allowed for independent translation of the RFP, providing a reliable indicator of successful vector delivery into the host cell, even in situations where the tagged IL-4 proteins might exhibit lower expression levels. To accomplish this, the vector pNonSynVP2-DsRed2 [81] containing a CMV immediate early promoter upstream of an IRES proceeding a DsRFP CDS was used. The coding sequence (828 bp) for monomeric GFP (muGFP) [79], flanked at its 5’ terminus by 75 bp encoding the vIL4 signal sequence and flanked at the 3’ terminus with 36 bp encoding a flexible linker, was synthesized commercially. Two constructs were generated using Gibson cloning in a 2 insert/1 vector scheme. The amplicons for the cIL-4 constructs were generated using primer Set 1: VTR muGFP FOR C-IL4/muGFP C-IL4REV cIL4 (muGFP amplicon); primer Set 2: C-IL4 MuGFP For/cIL4 Rev (cIL-4 amplicon); and primer Set 3: vtr cil4 for/vtrMuGFP REV C-IL4 (vector amplicon). The amplicon for the vIL4 constructs were generated using primer Set 4: VirIL4 For/muGFP vIL4 REV (muGFP amplicon); primers Set 5: vIL4 MuGFP For/VirIL4 Rev (vIL-4 amplicon); and primer Set 6: vtrVir IL4 for/vtr Vir IL rev (vector amplicon). For the sequences of the primers see Figure S1. Following in-fusion cloning the resulting cIL-4 and vIL-4 tagged constructs (Figure S2A) were transformed into NEB alpha electrocompetent E. coli and selected on LB agar plates containing kanamycin (50 μg/mL). Positive recombinants were identified through colony PCR and restriction enzyme digestion of mini-prep DNA.

### Transient expression and quantification of secreted GFP-tagged proteins

LMH cells were seeded into 6-well plates at a density of 8.0×10^5^ and transfected with 2.5 µg of GFP-tagged cIL-4 or GFP-tagged vIL-4 using TransIT (Mirus) according to the manufacturer’s instructions. Forty-eight hours post transfection the supernatants were harvested and centrifuged at 300×g for 10 minutes to remove cellular debris. Purified GFP standards (ranging from 10 ng/μL to 350 ng/μL) and diluted samples were pipetted (100 μL) into separate wells of a black, flat-bottomed 96-well plate. Fluorescence intensity was measured using a SpectraMax fluorometer at excitation and emission wavelengths of 488 nm and 507 nm, respectively. A standard curve was generated by plotting the average fluorescence intensity of each standard against its corresponding GFP concentration. The concentration of the secreted GFP-tagged protein in the original media was then determined by interpolating the sample fluorescence intensity on the standard curve.

### LPS stimulation of HD11 cells

HD 11 cells (2.0×10^5^) were grown in 96 wells overnight. When they reached ~85-90% con-fluency the culture media were replaced with fresh DMEM media to remove any immunostimulatory factors present in the conditioned media. To assess the direct effects of the GFP-tagged cytokines vIL-4 and cIL-4 on NO production, cells were preincubated with varying concentration (1000, 500, 250, 125 or 62.5 ng/ml) of either GFP-tagged vIL-4 or GFP-tagged cIL-4 for 4 hours. Following pre-incubation, LPS (Escherichia coli Serotype 0111:B4) 2.5 µg/ml was added to each well, and the cells were further stimulated for an additional 20 hours. Nitrite accumulation in the cell culture supernatant was measured as an indirect indicator of NO production using the Griess assay. Briefly, after the stimulation period, culture supernatants were transferred to a new plate and mixed with sulfanilamide and naphthylenediamine solutions. The plate was incubated for 30 minutes at room temperature, and nitrite concentration was determined by measuring the absorbance at 595 nm. A standard curve generated using sodium nitrite solutions was used to convert absorbance values into nitrite concentrations in the culture supernatants.

### Generation of vIL-4 null mutants

The sequence of the four vIL-4 exons were screened *in silico* using Geneious Prime software to identify optimal target sequences for Cas9 nuclease based on the scoring algorithms developed by Doench et al. [82]. Two sgRNAs ATCCTTAGGGAAGAACCCAGGT and ACCGTGCC-CTATAACAGTGGG with scores >0.7 were chosen and cloned in a human-codon optimized cas9 expression vector (VectorBuilder) under the control of U6 promoters (Figure S3). LMH cells were seeded in 6-well plates at a density of 1×10^5^ per well and incubated overnight at 39°C in 5% CO_2_ atmosphere. Cells at 90% confluency were transfected using TransIT (Mirus) with 7.5 µg of plasmid containing the sgRNAs and cas9 gene according to the manufacturer’s protocol. The transfected cells were incubated at 39°C in 5% CO_2_ for 12 hours. The cells were then infected with strain 1874C5 using either an MOI of 0.001 or 0.01. After absorption for one hour, the media was removed and DMEM containing 2% FBS and phosphonacetic acid (100 μg/ml) was added and the cells were incubated for an additional 12 hours. The media was again replaced with DMEM containing 2% FBS with 2% methylcellulose. Transfected/infected cells were incubated for 3 days, and 20 plaques were picked using a Evos microscope. Picked plaques, stored in a volume of 100-200 µl and frozen at −80°C, were subjected to three freeze/thaw cycles to release the virions.

### Plaque purification and PCR confirmation

To determine whether single plaques were isolated to homogeneity, a plaque PCR assay was employed. A single, isolated ILTV plaque from infected LMH cells overlayed with DMEM containing 2% methylcellulose was picked and suspended in complete DMEM medium. Virions were released after three freeze/thaw cycles at −80°C and 37°C, respectively. Following 5 rounds of plaque purification using limited dilutions (1:2) and expanding onto cells seeded in 24 well plates, the infected cells were trypsinized. To isolate the viral DNA, the cells were pelleted, washed with 1x PBS and lysed in 100 µl of 5mM Tris-HCl pH 8.0 at 95°C for 10 minutes. Proteinase K was added to a final concentration 0.3 µg/µl (1.5 µl of 20 mg/ml solution). After incubation at 55°C for 30 minutes, the reaction was stopped via heating at 95°C for 10 minutes and the cell debris removed by centrifugation. Five microliters were used in amplification reactions using LongAmp Taq DNA Polymerase (NEB) and primers: F1-4 AAGGGCCGCAATTAG-GTGGT and R1-4 AGCCCCGT TGACTTTAGCTT in a standard reaction with the following cycling conditions after an initial denaturation at 94°C for 2 min: 94°C for 30 sec, 58°C for 30 sec and 65°C for 1 min (40 cycles). The final extension was at 65°C for 1 min. Two clones (17 and A) that generated single amplicons of the expected size (654 bp) were selected for further confirmation. Sanger sequencing of the amplified region encompassing the targeted sequence verified the presence of the desired vIL-4 deletion.

### Replication kinetics

The replication of the ΔvIL4 (CK passage 2) as compared to its parental 1874C5 strain (CK passage 5) was evaluated in the LMH cell line. Briefly, LMH cells were seeded at 1.4×10^6^ cells in 2.5 ml of growth media (DMEM supplemented with 10% v/v fetal bovine serum) into 0.5% gelatin-treated six-well-plates (Cell BIND, Corning) and incubated at 37°C for 24 h. The cell monolayers were inoculated at an MOI of 0.002 with the ΔvIL4 or the parental 1874C5 strain. Viruses were adsorbed for 60 min at 39°C. After absorption the inoculum was removed and replaced with 2.5 ml of growth media supplemented with 2% fetal bovine serum. At 0 (immediately after media replacement), 24, 48, 72 and 96 hours post-incubation after replacing growth media, supernatant and cells from three independent wells (n=3) were separately collected and divided into two 1-ml aliquots for storage at −80°C. One of the aliquots was used to determine viral titer in LMH cells. Viral titers were expressed as the tissue culture infectious dose 50 perl ml (TCID_50_/ml) using the Reed and Meunch method [62], The second aliquot collected per well was utilized to extract total DNA and titrate viral genome copy numbers per sample.

### *Experimental design to assess* in vivo *expression of vIL-4*

A total of 60 specific pathogen free (SPF) eggs were obtained from a commercial source (Charles River Laboratories Inc., Wilmington, MA, USA) and incubated at 37.5°C and 55% relative humidity (RH) in a small-scale hatcher (Natureform Inc., Jacksonville, FL, USA) at the Poultry Diagnostic Research Center – University of Georgia. At 19 days of embryonation, eggs were candled and infertile eggs and nonviable embryos were removed. At hatch, 54 chickens were distributed in three groups of 18 and each group was housed in separate isolation units with filtered negative pressure air flow (6 chickens per unit). The chickens received water and a standard diet *ad libitum* during the duration of the experiment. At 18 days of age, two groups of chickens (n=18) were inoculated with either virulent strain 1874C5 or the CEO vaccine Laryngo-Vac (Zoetis, Animal Health). Viruses were administered at a dose of 10^3.8^ TCID_50_ in a 33 µL volume via eye drop. The third group of chickens (n=18) was mock inoculated in a similar fashion with cell culture media and served as the negative control (Mock). At 3-, 5-, and 7-days post-inoculation (p.i.), six chickens per group were humanely euthanized using CO_2_ inhalation. The upper and lower conjunctiva from both eyes were collected in 2 mL microcentrifuge tubes with 1 mL of RNA Later (Invitrogen, Waltham, MA, USA) and stored at −80°C for RNA extraction, cDNA synthesis, and library preparation for sequencing.

### Experimental design to assess parental 1874C5 and ΔvIL-4 pathogenicity

To evaluate whether the deletion of the vIL-4 gene changed the pathogenicity of the virulent 1874C5 strain, 24 days old specific pathogen free (SPF) chickens were inoculated with the parental 1874C5 or ΔvIL-4 strains via the intratracheal route. Briefly, seventy SPF eggs were obtained from a commercial source (ASV Bio Inc., Wilmington, MA, USA) and incubated at 37.5°C and 55% relative humidity (RH) in a small-scale hatcher (Natureform Inc., Jacksonville, FL, USA) at the Poultry Diagnostic Research Center – University of Georgia. At 19 days of embryonation, eggs were candled and infertile eggs and nonviable embryos were removed. At hatch, a total of 66 chickens were housed in isolation units with filtered negative pressure air flow. Six to seven chickens were housed per unit. Chickens received water and a standard diet *ad libitum* during the duration of the experiment. At 24 days of age, two groups of chickens (n=20 to 22) were inoculated with the parental 1874C5 or ΔvIL-4 strains. Chickens were inoculated with a 100 μl inoculum via the intratracheal route at a log_10_ 3.65 TCID_50_ dose of either viral strain. A third group of chickens (n=19) was mock-inoculated intratracheally with cell culture media. This group served as the negative control (Mock). From 2 to 7 days post inoculation (p.i.), clinical signs scores and percent survivability were documented. At 3- and 5-days p.i. tracheal swabs were collected from each group of chickens (Mock, parental 1875C5, and ΔvIL-4, n= 14 to 17) to evaluate virus genome load by real-time PCR. At four days p.i.five chickens that were not previously used to collect tracheal swabs were humanly sacrificed by CO_2_ inhalation from each group (Mock, parental 1875C5, and ΔvIL-4). The trachea was removed and an upper trachea segment (0.5 cm), caudal to the larynx, was collected from each chicken and fixed in 10% buffered formalin v/v, embedded in paraffin, sectioned (5 μm), and stained with hematoxylin and eosin (H&E) for examination of microscopic lesions and trachea epithelium changes induced after four days of ILTV inoculation. During the length of the experiment, chickens were given a standard feed diet, and water was provided *ad libitum*. All animal experiments described in this study were performed under the Animal Use Protocol A2018 06-009-Y2-A0 approved by the Animal Care and Use Committee (IACUC) in accordance with regulations of the Office of the Vice President for Research at the University of Georgia.

### Clinical signs

Clinical signs were evaluated for the categories of conjunctivitis, dyspnea, and lethargy. Each category was scored on a scale of 0 to 3, as previously described [6]. Briefly, the scoring of individual categories (of conjunctivitis, dyspnea, and lethargy) was considered 0 for normal; 0.5 or 1 for mild signs; 1.5 or 2 for moderate signs; and 2.5 or 3 for severe signs. The total clinical score per chicken was calculated based on the sum of the three categories scores. The sum of scores for individual clinical sign categories (total clinical signs) per chicken, and the average clinical sign total score per group per time point was calculated.

### Trachea microscopic lesions

Upper trachea sections collected at four days p.i. from Mock, parental 1874C5 and ΔvIL-4-inoculated chickens were microscopically examined. Each trachea section received a score to reflect the level of microscopic lesions induced by ILTV infection. Lesions were scored according to the criteria previously described by Guy et al [37]. Briefly, a score scale from 0 to 5 was utilized. Score 0: normal epithelium. Score 1: normal epithelium with mild to moderate lymphocytic infiltration and absence of syncytia with intranuclear inclusion bodies. Score 2: normal epithelium with mild to moderate lymphocytic infiltration and few foci of syncytia with intranuclear inclusion bodies. The latter are the two characteristic lesions that define ILTV lytic replication. Score 3: affected epithelium shows moderate to marked hyperemia, lymphocytic infiltration, and numerous syncytia with intranuclear inclusion bodies. Score 4: mucosa shows areas without epithelium and syncytia with intranuclear inclusion bodies sometimes present. Score 5: Mucosa has no residual epithelium and occasional presence of syncytia and intranuclear inclusion bodies.

Also, the amount of mucosa epithelium and submucosa remaining in the trachea rings collected from Parent 1874C5 IT and ΔvIL-4 IT groups of chickens was assessed microscopically. Briefly, each trachea ring (n=5 per group) was divided digitally in six segments (1 to 6 clockwise) and in each segment (n=30 per group) was scored. A score of 0 was conferred to trachea segments where only a thin layer of submucosa distal to the trachea cartilage remained. A score of 1 was conferred to trachea segments with expanded submucosa, due to increased infiltration, but without any mucosal epithelium remaining. A score of 2 was conferred to trachea segments where submucosa with a distal thin layer of mucosal epithelium remained in place after four days post inoculation. The number of trachea segments categorized as score of 0, 1, or 2 for chickens inoculated with the parental 1874C5 or the ΔvIL-4 strain are presented.

### DNA extraction

Tracheal swabs were placed in 2 ml tubes with 1×PBS. All samples were stored at −80°C until processed. Total DNA from feather pulps and tracheas were extracted using the MagaZorb DNA mini-prep kit (Promega, Madison, WI), following the manufacturer’s recommendations with a few modifications. Briefly, using a 96-well plate, 7 μl of proteinase K (PK) solution was loaded into each well, followed by 70 μl of sample and 50 μl of lysis buffer per well. The plate was then incubated at 56°C for 10 min, 10 μl of magnetic beads and 125 μl of binding buffer were added per well, followed by rotating the plate for 10 min at room temperature. The supernatant and magnetic beads were separated using a magnetic stand and the supernatant was discarded. Finally, using 250 μl of washing buffer, the beads were washed twice, and DNA was eluted from the beads with 100 μl of elution buffer after rotating the plate for 10 min at room temperature.

### *RNA extraction, cDNA synthesis, and RNA-Seq for* in vivo *expression*

Conjunctivas were thawed and transferred into lysing bead matrix tubes (MP Biomedicals, Santa Ana, CA) containing 900 μL of Qiazol (Qiagen, Germantown, MD, USA), incubated for 30 minutes and homogenized using the FastPrep-24 5G (MP Biomedical, Santa Ana, CA, USA) at 6.0 m/s for 20 seconds. Total RNA from conjunctiva tissues was extracted using the RNeasy Plus universal kit (Qiagen, Germantown, MD, USA) according to the manufacturer’s protocol. RNA concentrations and purity were determined with the Nanodrop 2000c spectrophotometer (ThermoFisher Scientific, Waltham, MA, USA). cDNA was synthesized from 1 µg of total RNA using the RT2 First Strand Kit (Qiagen, Germantown, MD, USA), following the manufacturer’s guidelines. Stranded sequencing libraries were prepared from ribo-depleted RNA preparations using KAPA Stranded mRNA kit and sequenced on an Illumina NextSeq 2000 instrument using a high output flow cell at the Georgia Genomics and Bioinformatic Core at the University of Georgia.

### In vivo *RNA-Seq data analysis*

Raw FASTQ files were trimmed using Trim Galore (https://github.com/FelixKrueger/TrimGalore) (`--2colour 8 --stringency 4 --length 40 --trim-n --max_n 5`). Kallisto v0.50.1 was used for transcript quantification [83]. A reference index was generated using `kallisto index` on the combined bGalGal1.mat.broiler.GRCg7b + virus transcript sequences. For this purpose, a locally updated annotation of the 1874C5 genome was used, based on a corrected genome sequence. These files in GFF3 and FASTA format are available in the git repository for this project (see Data Availability) and upon request. Quantification was performed with the following non-default settings: `--rf-stranded`. TPM values were merged for all replicates and re-normalized using only viral transcripts.

### Protein extraction and LC-MS/MS

Primary chicken kidney cells propagated in F75 flasks were infected with ILTV virulent strain 1874C5 or vaccine strain LaryngoVac (moi of 0.001), or mock-infected with culture media. At 72 hr post-infection, the supernatants were removed and the cells harvested via scraping. Following centrifugation, the aspirated cells were frozen at −80°C. Three replicates of each were shipped to the University of Illinois at Urbana-Champaign Proteomics Core Facility as frozen pellets.

Cell pellets were subsequently lysed in a buffer containing 6M guanidine HCl, 10 mM tris(2-carboxyethyl)phosphine HCL, and 40 mM 2-chloroacetamide and then boiled to promote reduction and alkylation of disulfide bonds, as previously described [84]. The samples were cleared of debris by centrifugation and the pH was adjusted to 8 with 100 mM triethylammonium bicarbonate. Protein amounts were determined by BCA assay (Pierce, Rockford, IL) before sequential proteolytic digestion by LysC (1:100 w/w enzyme: substrate; Wako Chemicals, Richmond, VA) for 4h at 30°C and trypsin (1:50 w/w; Pierce, Rockland, IL) overnight at 37°C. Peptide samples were desalted using StageTips and dried in a vacuum centrifuge. Peptide digests were analyzed using a Thermo UltiMate 3000 UHPLC system coupled to a high resolution Thermo Q Exactive HF-X mass spectrometer. Peptides were separated by reverse-phase chromatography using a 50 cm Acclaim PepMap 100 C18 column maintained at 60°C with mobile phases of 0.1% formic acid (A) and 0.1% formic acid in 80% acetonitrile (B). A two-step linear gradient from 5% B to 35% B over the course of 110 min and 35% B to 50% B over 10 min was employed for peptide separation, followed by additional steps for column washing and equilibration. The MS was operated in a data-dependent manner in which precursor scans from 350 to 1500 m/z (120,000 resolution) were followed by higher-energy collisional dissociation (HCD; 30 NCE) of the 20 most abundant ions. MS2 scans were acquired at a resolution of 15,000 with a precursor isolation window of 1.0 m/z and a dynamic exclusion window of 60 seconds.

### Proteogenomic analysis

A proteogenomic search database containing the host (bGalGal1.mat.broiler.GRCg7b) proteome, viral known proteome, viral six-frame translation, viral novel splice junction peptides, and viral potential alternative translation start peptides, along with potential contaminants and reversed decoy sequences, was built as previously described for GaHV-2 [85]. Database searching using Comet and MS-GF+, as well as post-processing with Percolator, were carried out exactly as described in the same work.

Label-free peptide quantification was performed using FlashLFQ v2.0.0 [86] with match-between run (MBR), inter-sample normalization, and requiring within-condition PSM for MBR (`--nor true --mbr --rmc`). Peptide intensities were used to calculate protein iBAQ values by dividing summed peptide intensities for each protein by the number of theoretical fully tryptic peptides length 6-40 in the protein. Intensities for peptides shared between proteins were divided evenly between the group members. Relative iBAQ (riBAQ) [87] was calculated within each replicate as the protein iBAQ divided by the sum of all iBAQ values for the replicate, considering only viral proteins.

### Re-analysis of LJS09 sequencing

RNA-Seq reads originally published by Wang et al. [21] of treatment of chick embryo liver cells with ILTV strain LJS09 or mock were downloaded from the NCBI Sequence Read Archive (accessions SRR5515169–SRR5515172 and SRR5515181–SRR5515184). These reads were mapped directly against the combined bGalGal1.mat.broiler.GRCg7b + virus genome reference using HISAT2 v2.2.1 [88] (defaults) and converted to bedgraph coverage format for visualization using bedtools v2.31.1 (`genomecov -split -ibam <IN> -bg`) [89].

### Duplex real-time PCR

ILTV genomes were quantified from LMH infected cultures and from tracheal swabs collected at days 3 and 5 p.i. using a duplex real-time PCR. The ILTV genomes were amplified with primers that target the UL44 ILTV gene, and chicken genome was detected with primers that target the chicken α2 collagen gene as previously described [90]. The relative amount of viral DNA per sample was calculated as the log_10_ 2-ΔΔC^t^ as previously described [91].

### *Statistical analysis of replication and* in vivo *results*

The Shapiro–Wilk test was used to test the normality of the data. ILTV genome load data were analyzed using one-way ANOVA followed by Tukey’s multiple comparisons test and with the non-parametric Mann-Whitney test. Clinical signs and trachea histopathological microscopic scores were evaluated using the Kruskal–Wallis test followed by the Dunn’s multiple comparisons test and with the non-parametric Mann-Whitney test. All statistical analyses were performed using the GraphPad Prism software 10 (GraphPad Software, Inc., La Jolla, CA). All reported p-values were based on two-sided comparisons. Statistical differences were defined as significant when they reached p<0.05.

### Manuscript preparation

The manuscript was prepared using RMarkdown with heavy use of the ggplot2 library [92] for generating figures. Figure 1 was generated using the Gviz R package v1.46.1 [93]. Pymol [94] was used for visualization of 3D modeling. The ggtree [95] R package was used for tree visualization. All supplementary code used for data format manipulation and processing is available from the manuscript git repository (see *Data Availability*).

## Supplementary materials

**Figure S1:**
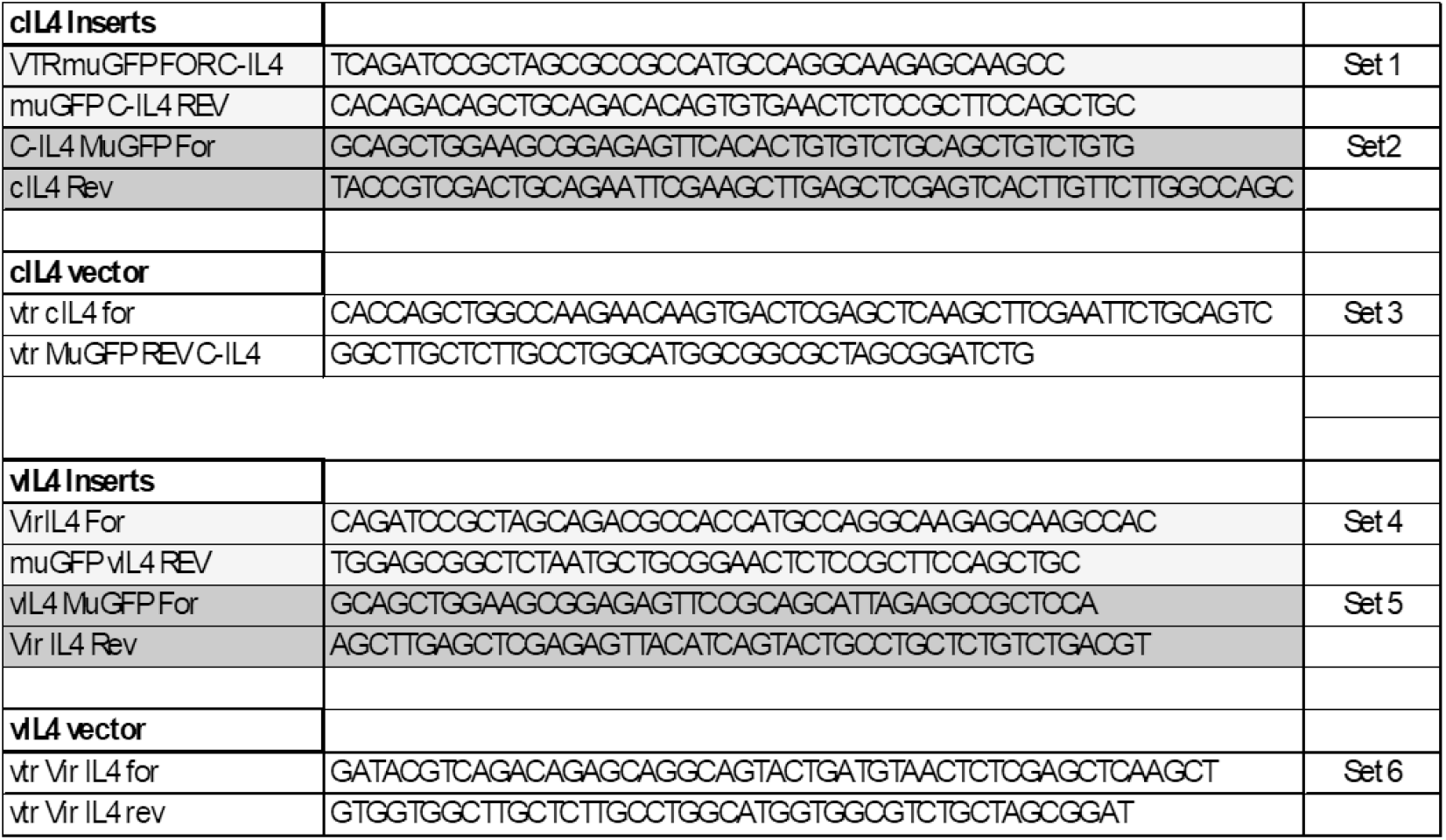
List of primers used to generate PCR amplicons, which were assembled to create N-terminal tagged versions of cIL-4 and vIL-4.

**Figure S2:**
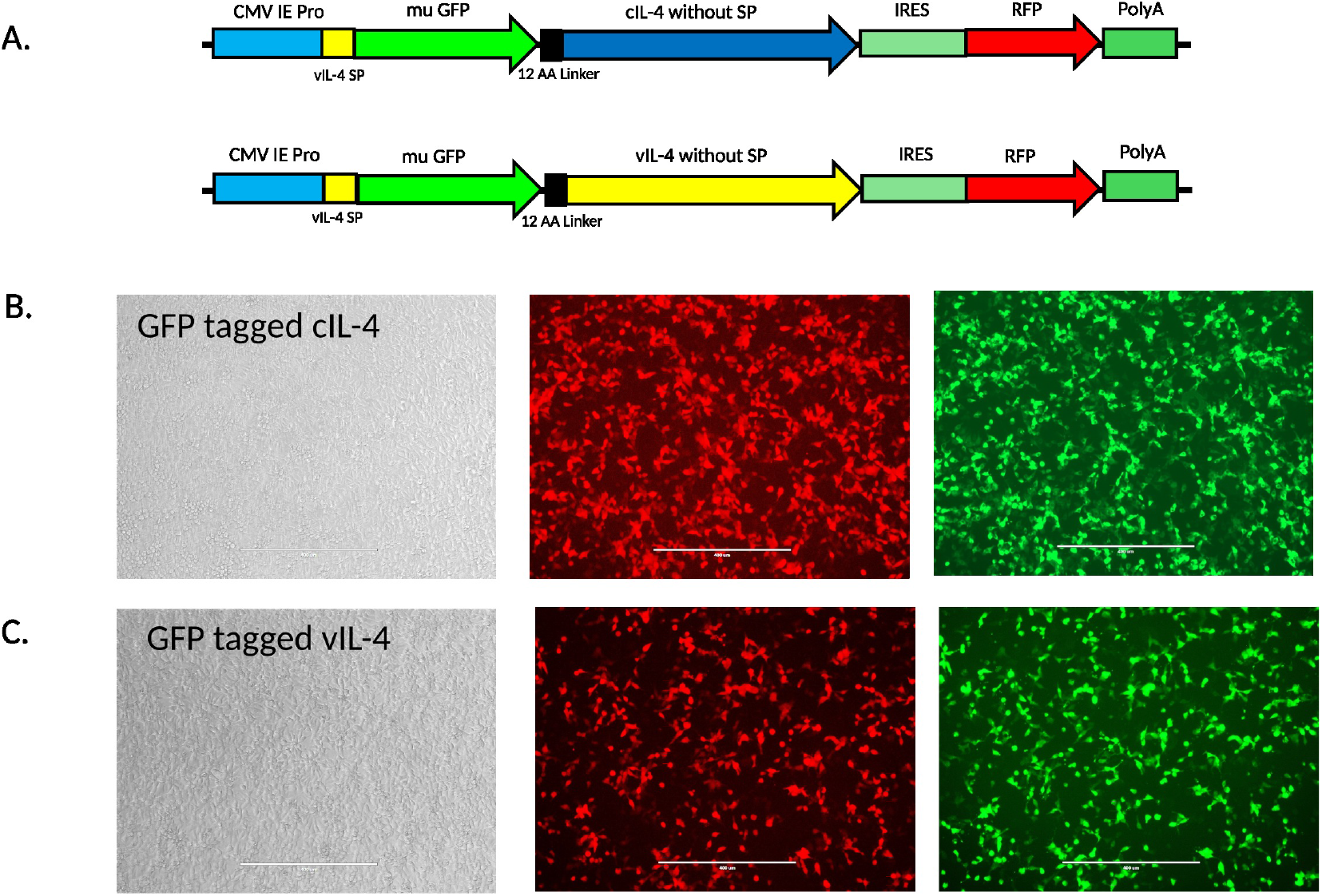
GFP-tagged cIL-4 and vIL-4. (A) Diagram of the plasmid constructs expressing GFP-tagged cIL-4 and vIL-4 proteins. Fluorescent microscopy of LMH cells transfected with (B) GFP-cIL-4 and (C) GFP-vIL4. Magnification is 100×.

**Figure S3:**
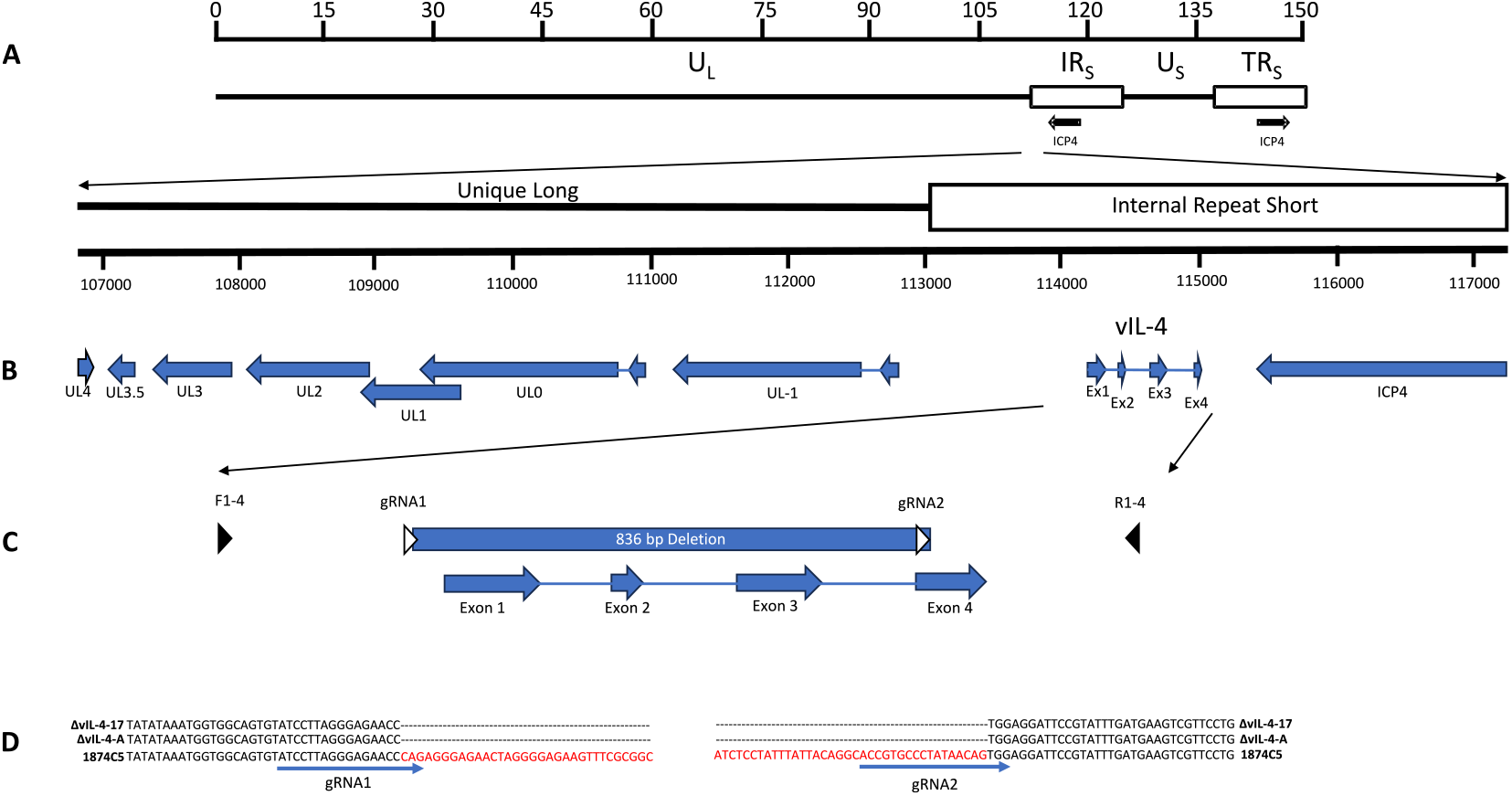
Generation of a vIL-4 deletion mutant. (A) Schematic of the ILTV D-type genome, showing the unique long (UL) and unique short (US) regions flanked by inverted repeats (internal repeat short [IRS] and terminal repeat short [TRS]). (B) Gene organization at the UL/TRS junction, indicating gene direction. (C) The vIL-4 locus, with exons (arrows connected by lines) and their corresponding coordinates based on GenBank entry JN542533: Exon 1 (114,156–114,302), Exon 2 (114,422–114,468), Exon 3 (114,629–114,768), and Exon 4 (114,919–115,028). White triangles indicate the guide RNA (gRNA) binding sites. Black triangles denote the locations of forward (F1-4, TAAGGCCGCAATTAGGTGCT; 113,787–113,806) and reverse (R1-4, AGCCC-CGTTGACTTTAGCTT; 115,278–115,259) oligonucleotide primers used for amplicon generation. (D) Sanger sequencing alignment of amplicons from two plaque-purified ΔvIL-4 mutants (#17 and A). Red nucleotides indicate the deleted sequences adjacent to the gRNA binding sites.

**Figure S4:**
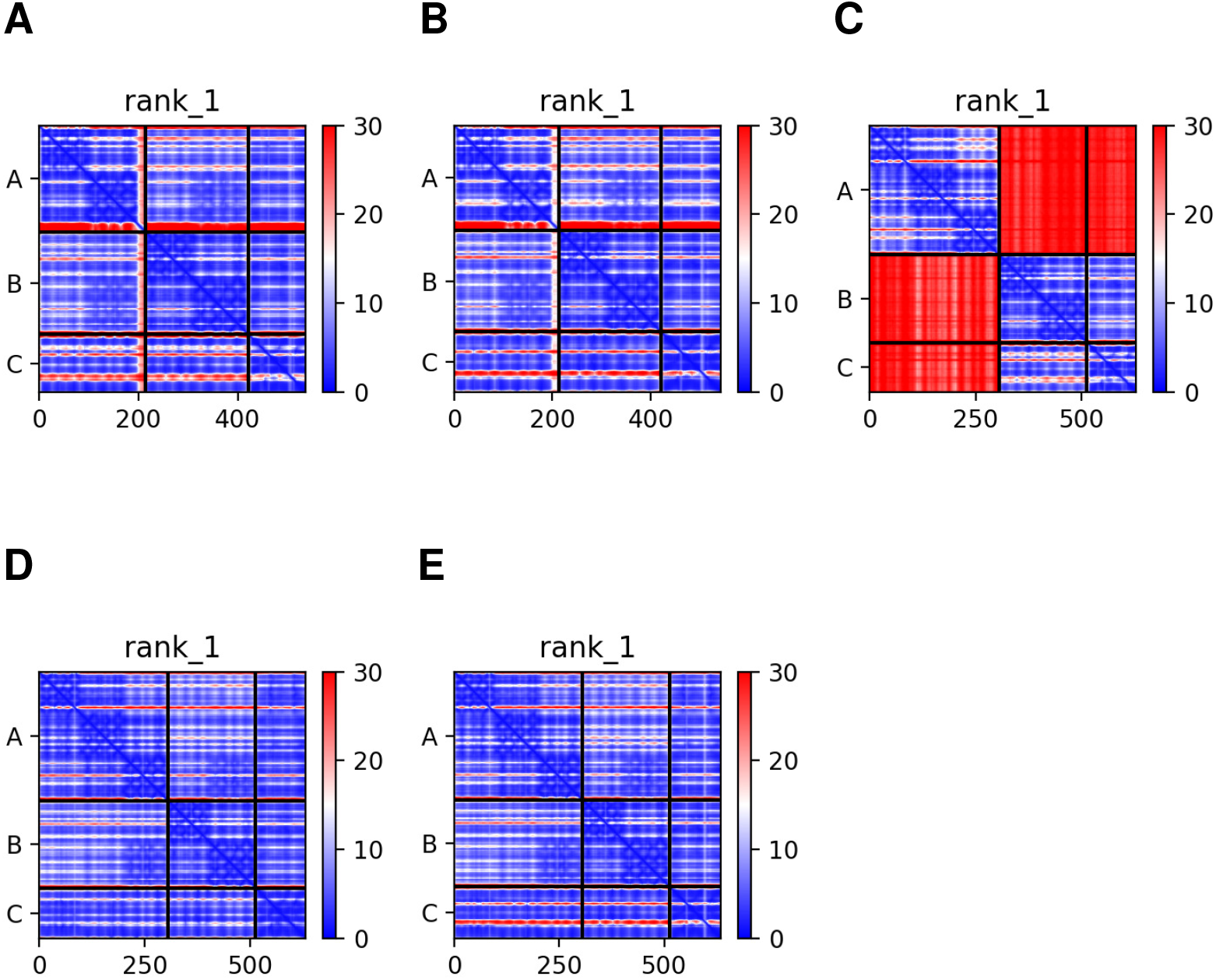
Predicted aligned error (PAE) plots of AlphaFold multimeric models for (A) gamma-c + IL-4Ra + cIL-4; (B) gamma-c + IL-4Ra + vIL-4; (C) IL-13Ra1 + IL-4Ra + cIL-4; (D) IL-13Ra1 + IL-4Ra + cIL-13; (E) IL-13Ra1 + IL-4Ra + vIL-4.

**Figure S5:**
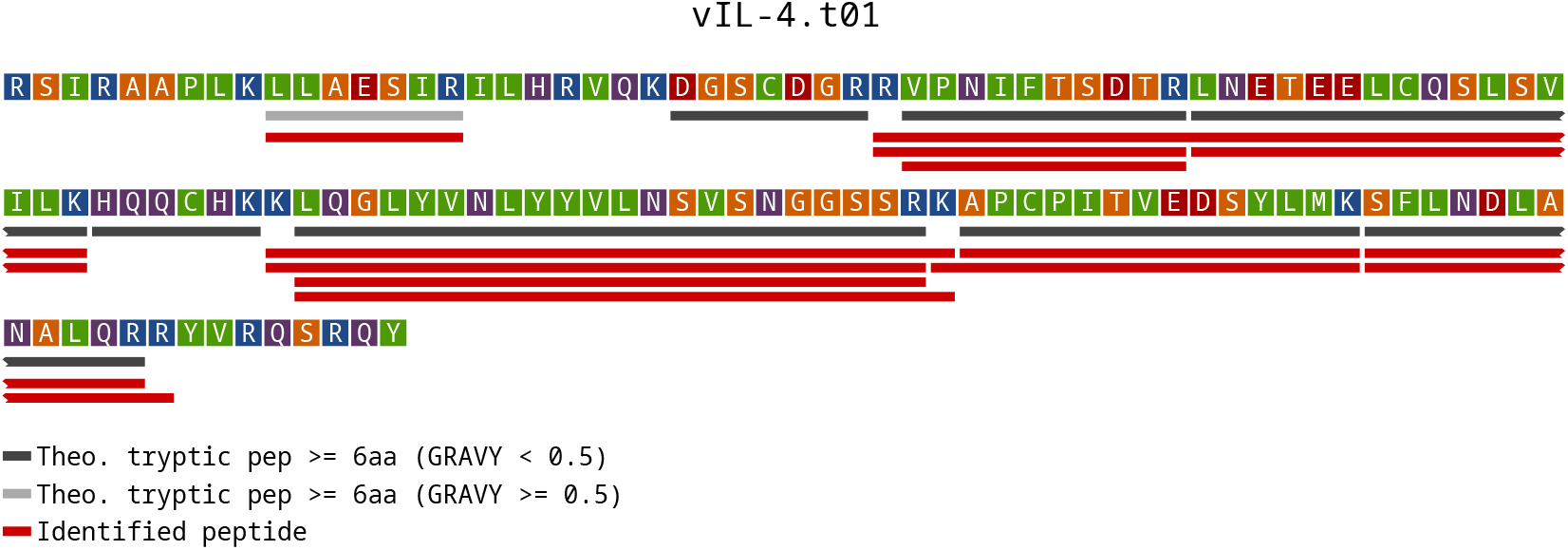
Map of identified and theoretical trypic peptides within the vIL-4 protein. The putative leader sequence has been removed. Red bars are identified peptides (1% FDR) including peptides with missed cleavages. Dark gray bars indicate theoretical tryptic peptides with GRAVY (GRand AVerage of hydropathY) score < 0.5. Light gray bars indicate theoretical tryptic peptides with GRAVY score >= 0.5 (more hydrophobic and thus possibly more difficult to detect). Tryptic peptides shorter than 6 aa were not searched and are not shown.

**Figure S6:**
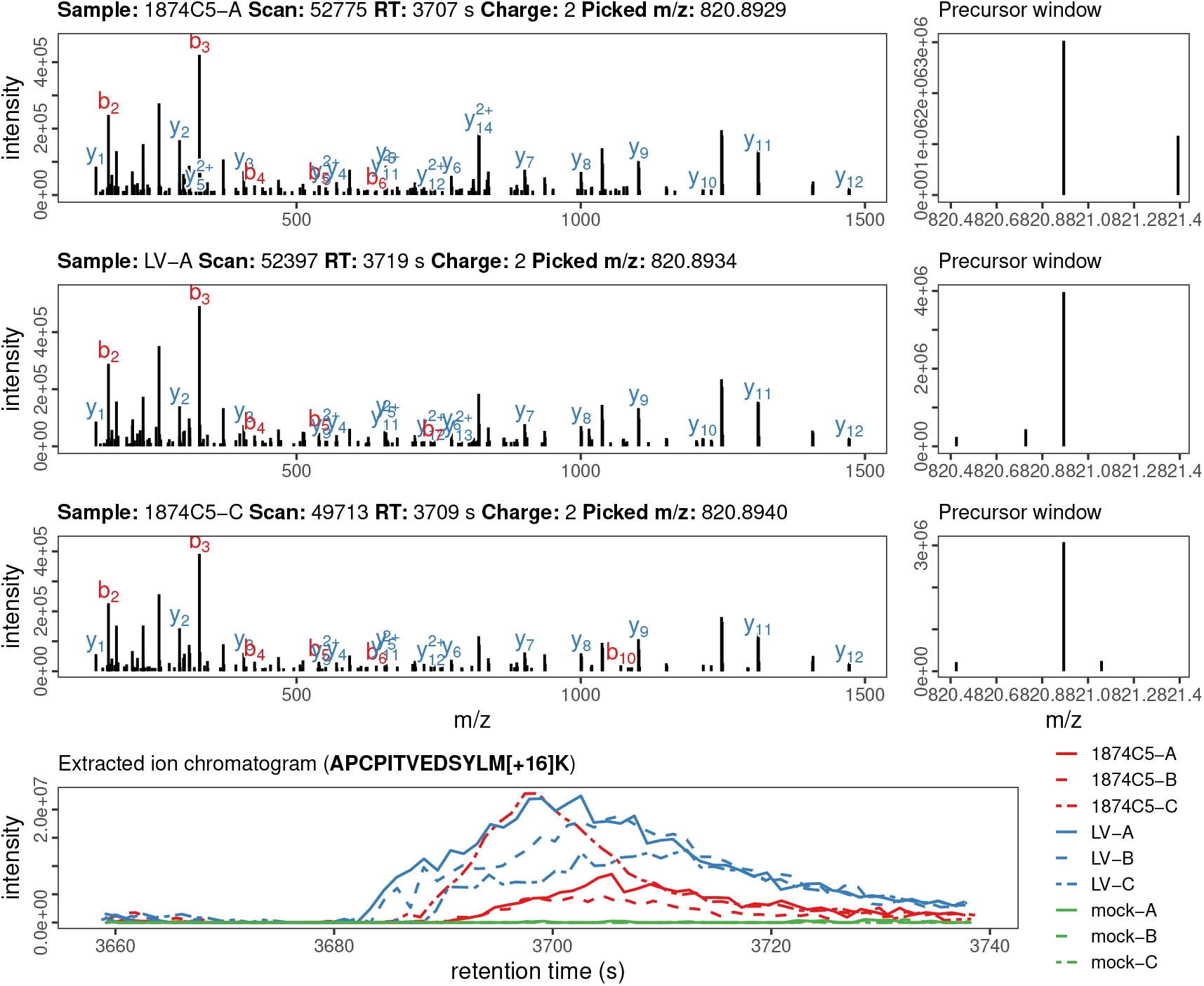
QC data for vIL-4 peptide APCPITVEDSYLM[+16]K identified by LC-MS/MS. Top rows show up to three MS2 spectra matched at 1% FDR, with matching y-b ion series labeled in blue and red. On the right of each row is the MS1 precursor window used for fragmentation. The bottom row shows an extracted ion chromatogram (XIC) for the peptide precursor mass over the matching retention time window for all nine samples (1875C5, LaryngoVac, and mock). True viral peptides should have XIC elution peaks in infected samples but not mock.

**Figure S7:**
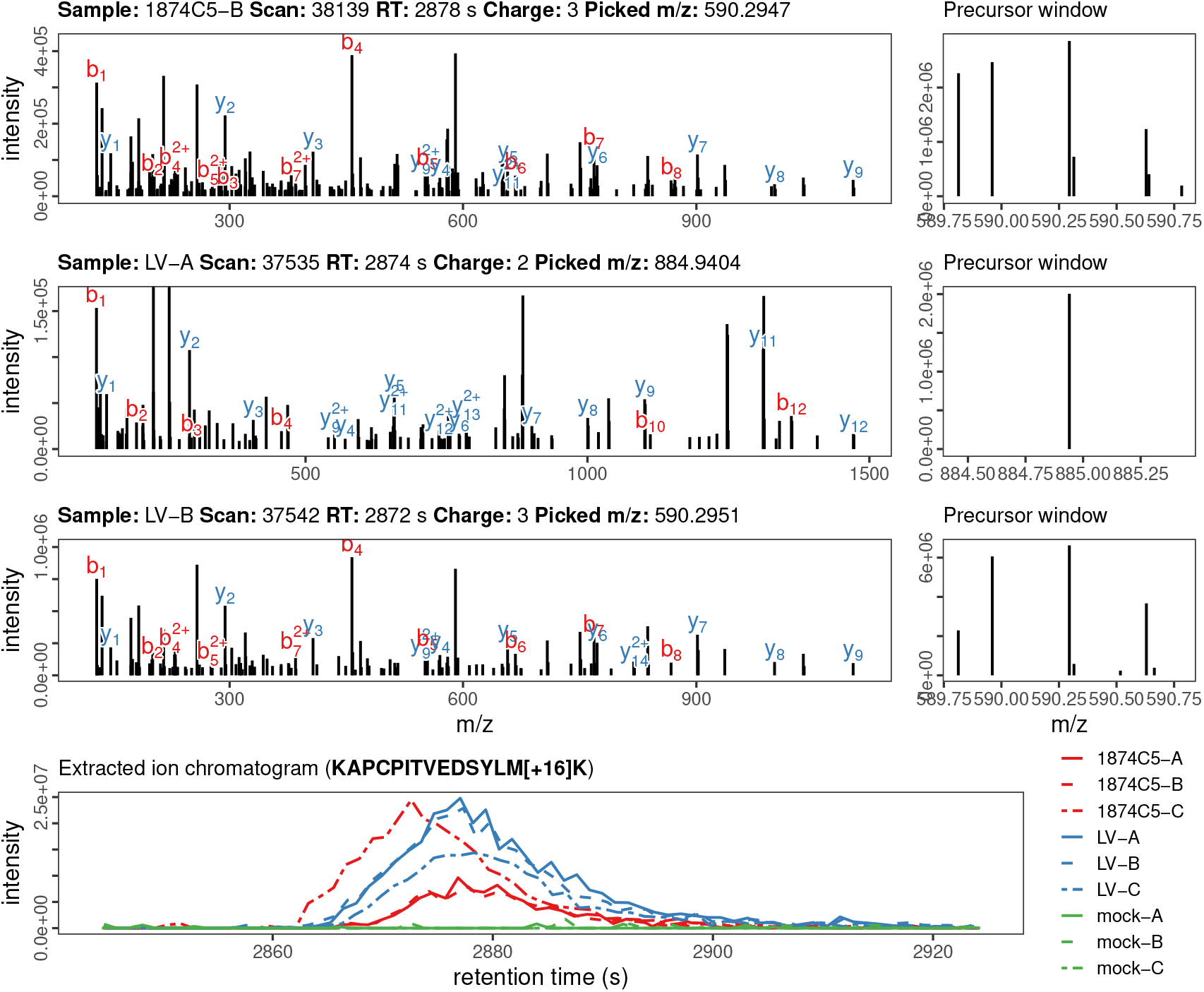
QC data for vIL-4 peptide KAPCPITVEDSYLM[+16]K identified by LC-MS/MS. Top rows show up to three MS2 spectra matched at 1% FDR, with matching y-b ion series labeled in blue and red. On the right of each row is the MS1 precursor window used for fragmentation. The bottom row shows an extracted ion chromatogram (XIC) for the peptide precursor mass over the matching retention time window for all nine samples (1875C5, LaryngoVac, and mock). True viral peptides should have XIC elution peaks in infected samples but not mock.

**Figure S8:**
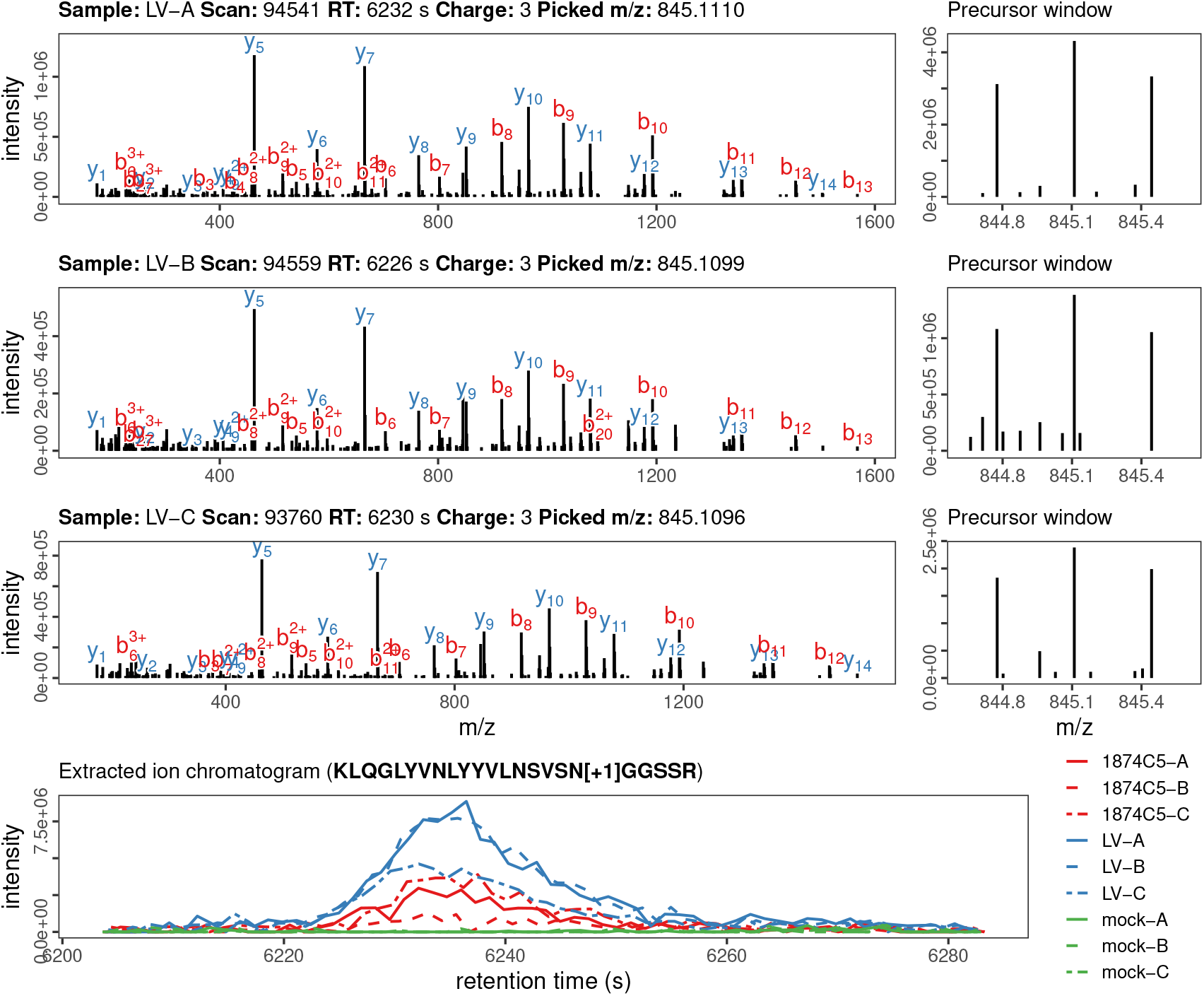
QC data for vIL-4 peptide KLQGLYVNLYYVLNSVSN[+1]GGSSR identified by LC-MS/MS. Top rows show up to three MS2 spectra matched at 1% FDR, with matching y-b ion series labeled in blue and red. On the right of each row is the MS1 precursor window used for fragmentation. The bottom row shows an extracted ion chromatogram (XIC) for the peptide precursor mass over the matching retention time window for all nine samples (1875C5, LaryngoVac, and mock). True viral peptides should have XIC elution peaks in infected samples but not mock.

**Figure S9:**
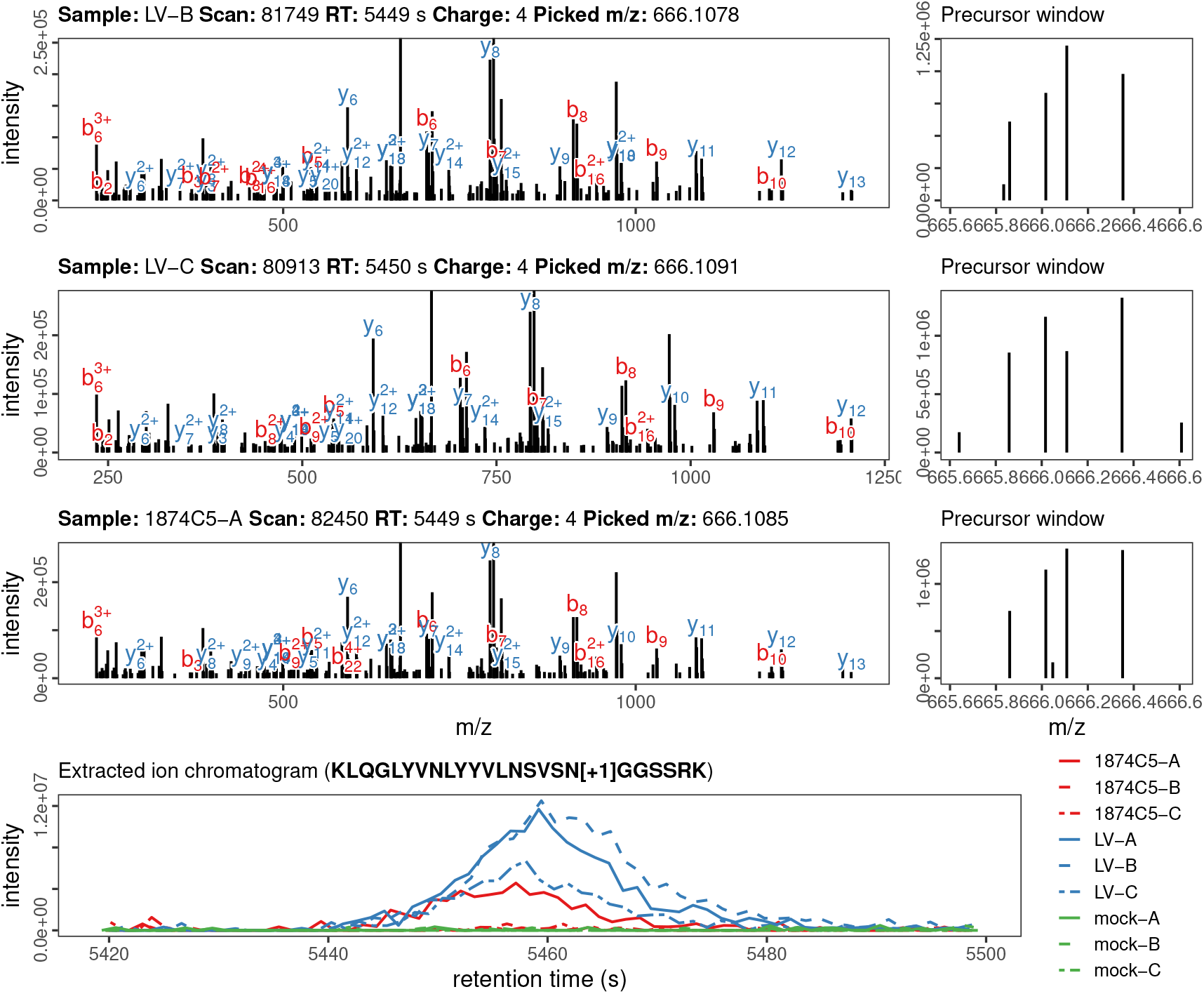
QC data for vIL-4 peptide KLQGLYVNLYYVLNSVSN[+1]GGSSRK identified by LC-MS/MS. Top rows show up to three MS2 spectra matched at 1% FDR, with matching y-b ion series labeled in blue and red. On the right of each row is the MS1 precursor window used for fragmentation. The bottom row shows an extracted ion chromatogram (XIC) for the peptide precursor mass over the matching retention time window for all nine samples (1875C5, LaryngoVac, and mock). True viral peptides should have XIC elution peaks in infected samples but not mock.

**Figure S10:**
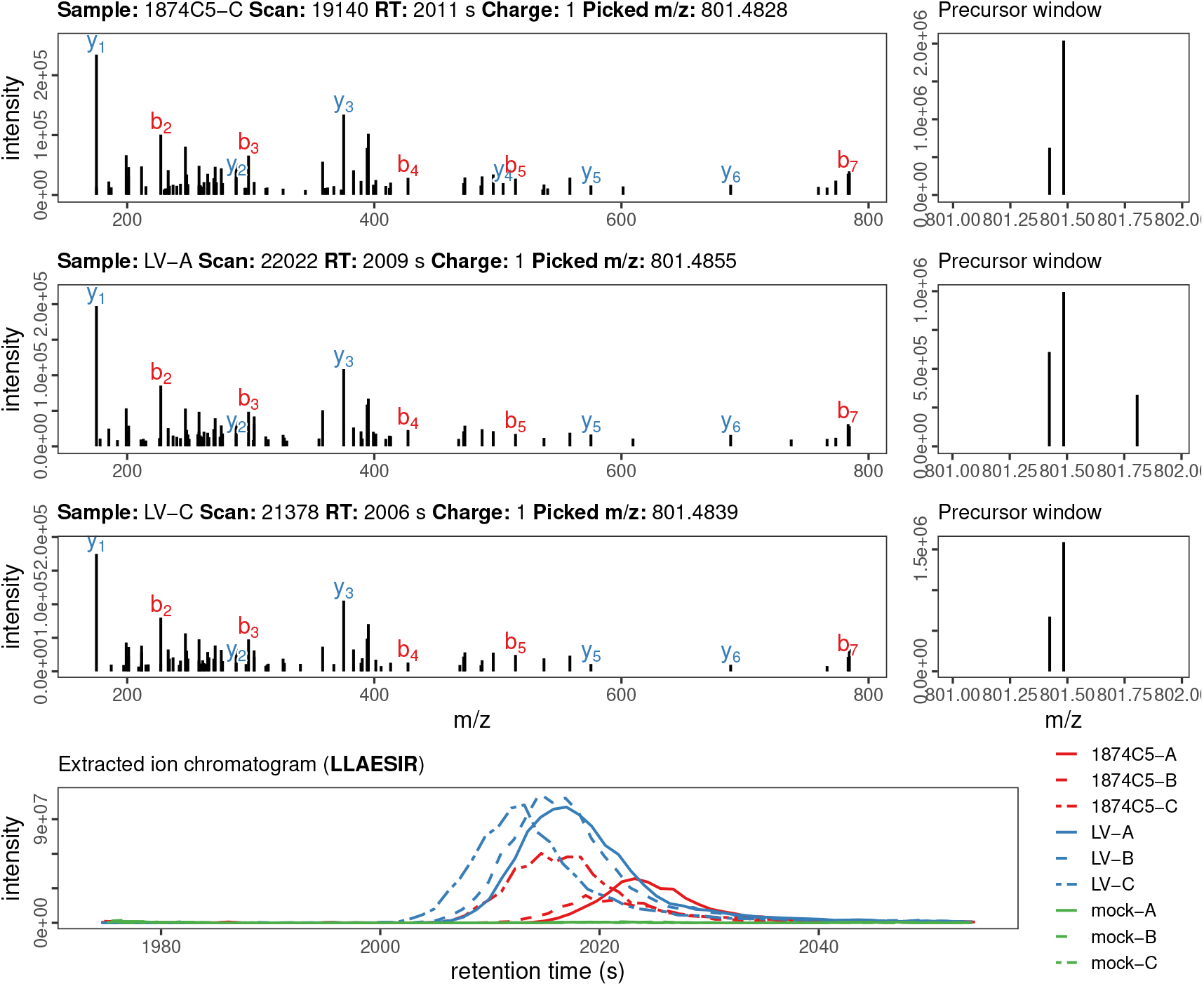
QC data for vIL-4 peptide LLAESIR identified by LC-MS/MS. Top rows show up to three MS2 spectra matched at 1% FDR, with matching y-b ion series labeled in blue and red. On the right of each row is the MS1 precursor window used for fragmentation. The bottom row shows an extracted ion chromatogram (XIC) for the peptide precursor mass over the matching retention time window for all nine samples (1875C5, LaryngoVac, and mock). True viral peptides should have XIC elution peaks in infected samples but not mock.

**Figure S11:**
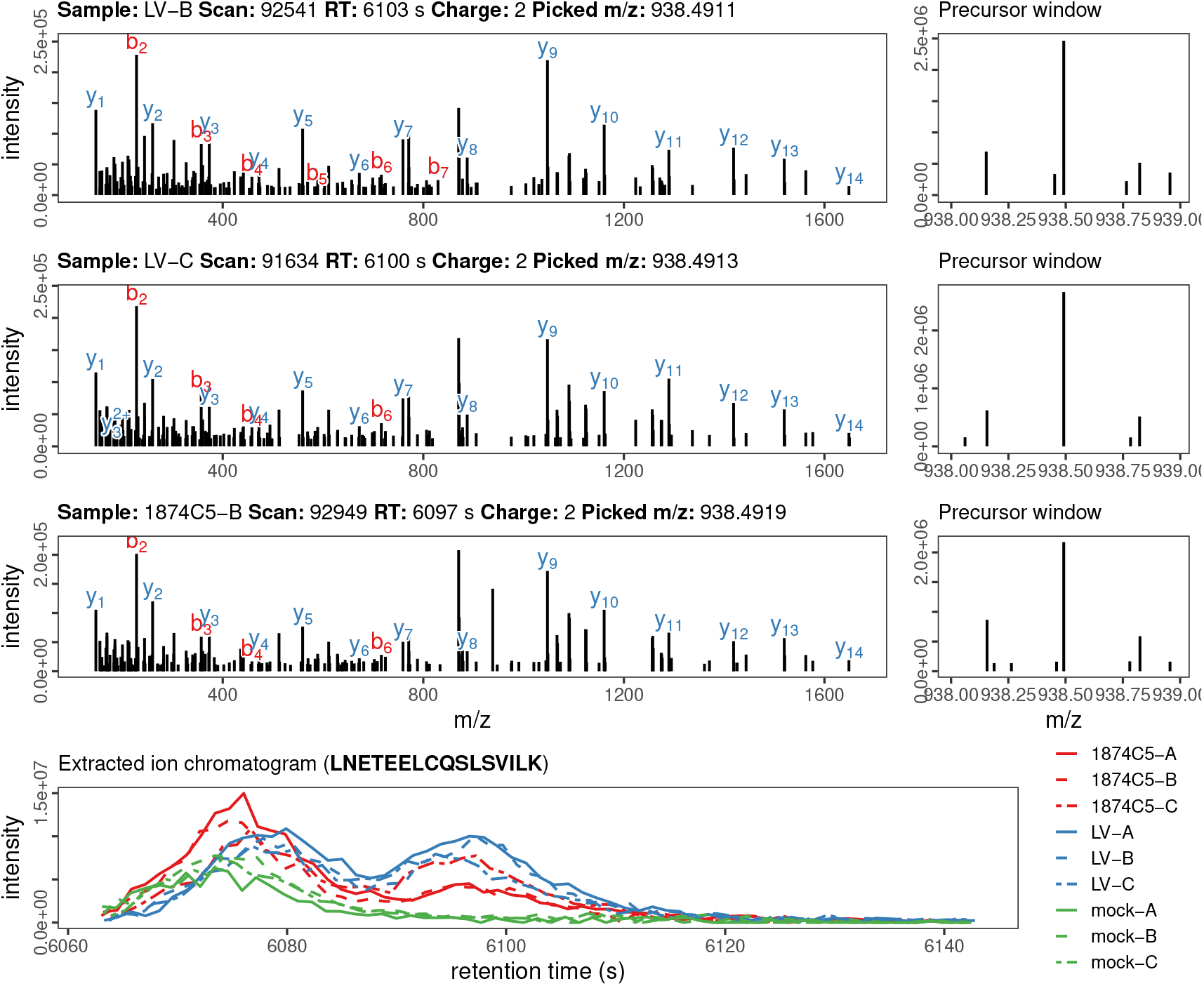
QC data for vIL-4 peptide LNETEELCQSLSVILK identified by LC-MS/MS. Top rows show up to three MS2 spectra matched at 1% FDR, with matching y-b ion series labeled in blue and red. On the right of each row is the MS1 precursor window used for fragmentation. The bottom row shows an extracted ion chromatogram (XIC) for the peptide precursor mass over the matching retention time window for all nine samples (1875C5, LaryngoVac, and mock). True viral peptides should have XIC elution peaks in infected samples but not mock.

**Figure S12:**
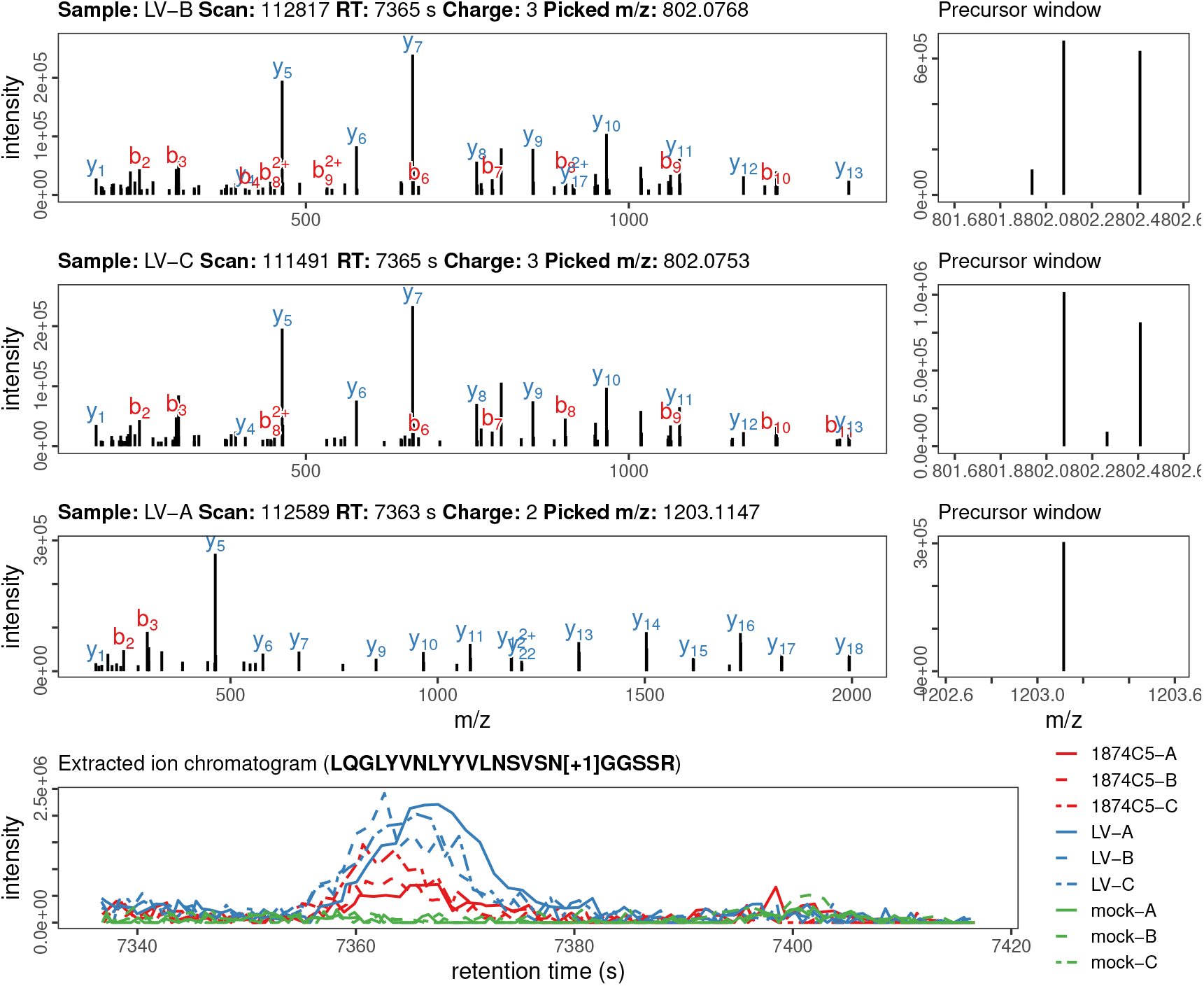
QC data for vIL-4 peptide LQGLYVNLYYVLNSVSN[+1]GGSSR identified by LC-MS/MS. Top rows show up to three MS2 spectra matched at 1% FDR, with matching y-b ion series labeled in blue and red. On the right of each row is the MS1 precursor window used for fragmentation. The bottom row shows an extracted ion chromatogram (XIC) for the peptide precursor mass over the matching retention time window for all nine samples (1875C5, LaryngoVac, and mock). True viral peptides should have XIC elution peaks in infected samples but not mock.

**Figure S13:**
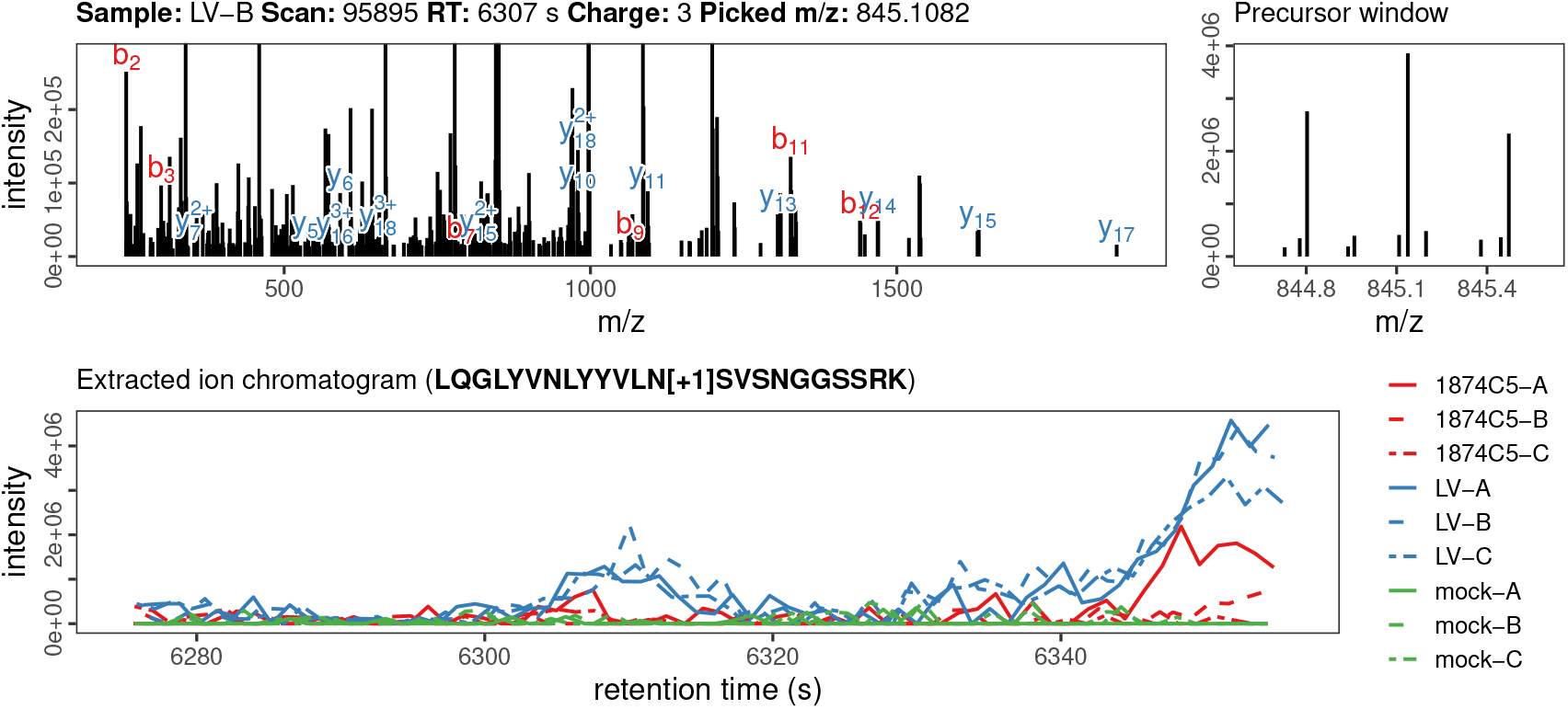
QC data for vIL-4 peptide LQGLYVNLYYVLN[+1]SVSNGGSSRK identified by LC-MS/MS. Top rows show up to three MS2 spectra matched at 1% FDR, with matching y-b ion series labeled in blue and red. On the right of each row is the MS1 precursor window used for fragmentation. The bottom row shows an extracted ion chromatogram (XIC) for the peptide precursor mass over the matching retention time window for all nine samples (1875C5, LaryngoVac, and mock). True viral peptides should have XIC elution peaks in infected samples but not mock.

**Figure S14:**
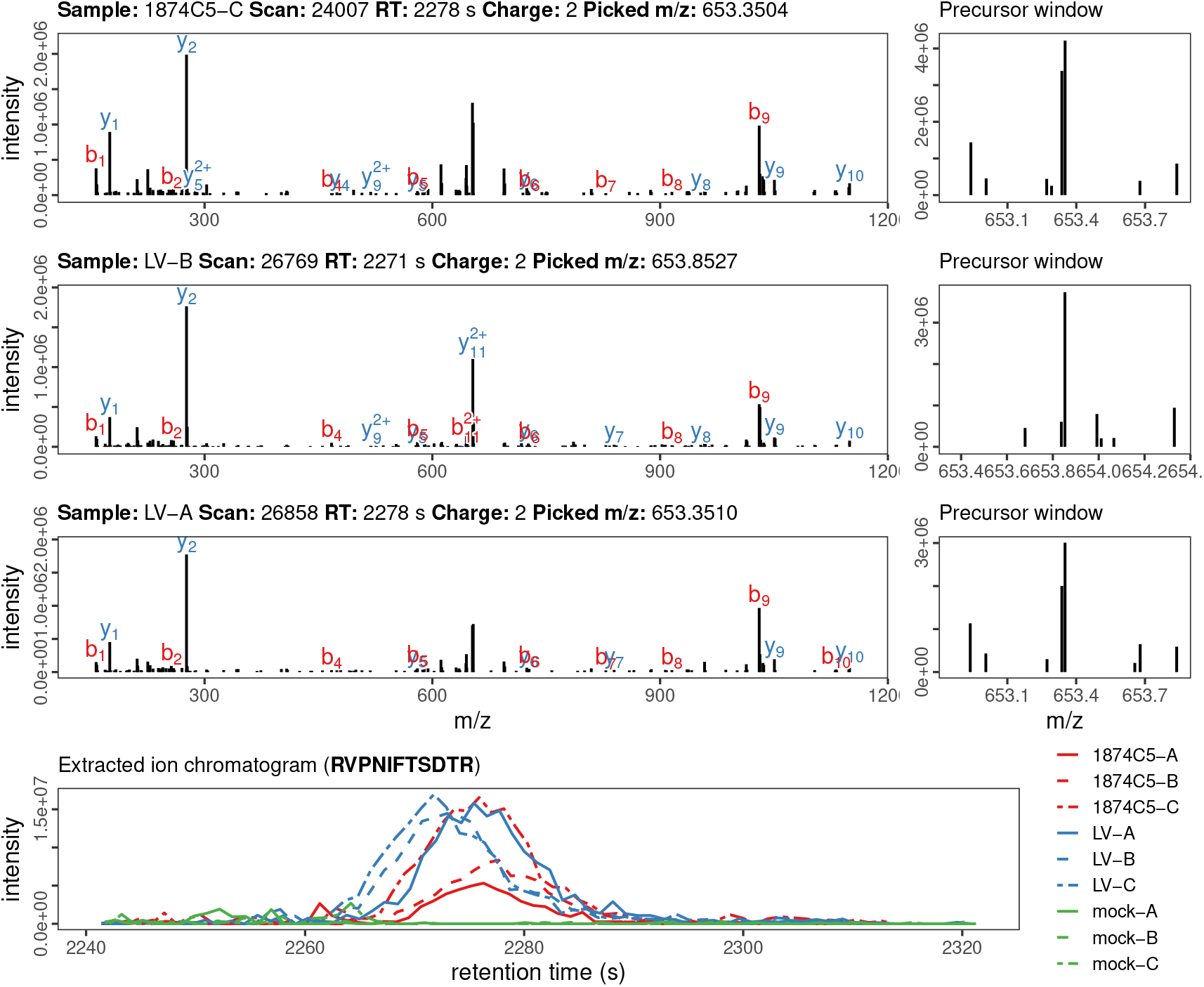
QC data for vIL-4 peptide RVPNIFTSDTR identified by LC-MS/MS. Top rows show up to three MS2 spectra matched at 1% FDR, with matching y-b ion series labeled in blue and red. On the right of each row is the MS1 precursor window used for fragmentation. The bottom row shows an extracted ion chromatogram (XIC) for the peptide precursor mass over the matching retention time window for all nine samples (1875C5, LaryngoVac, and mock). True viral peptides should have XIC elution peaks in infected samples but not mock.

**Figure S15:**
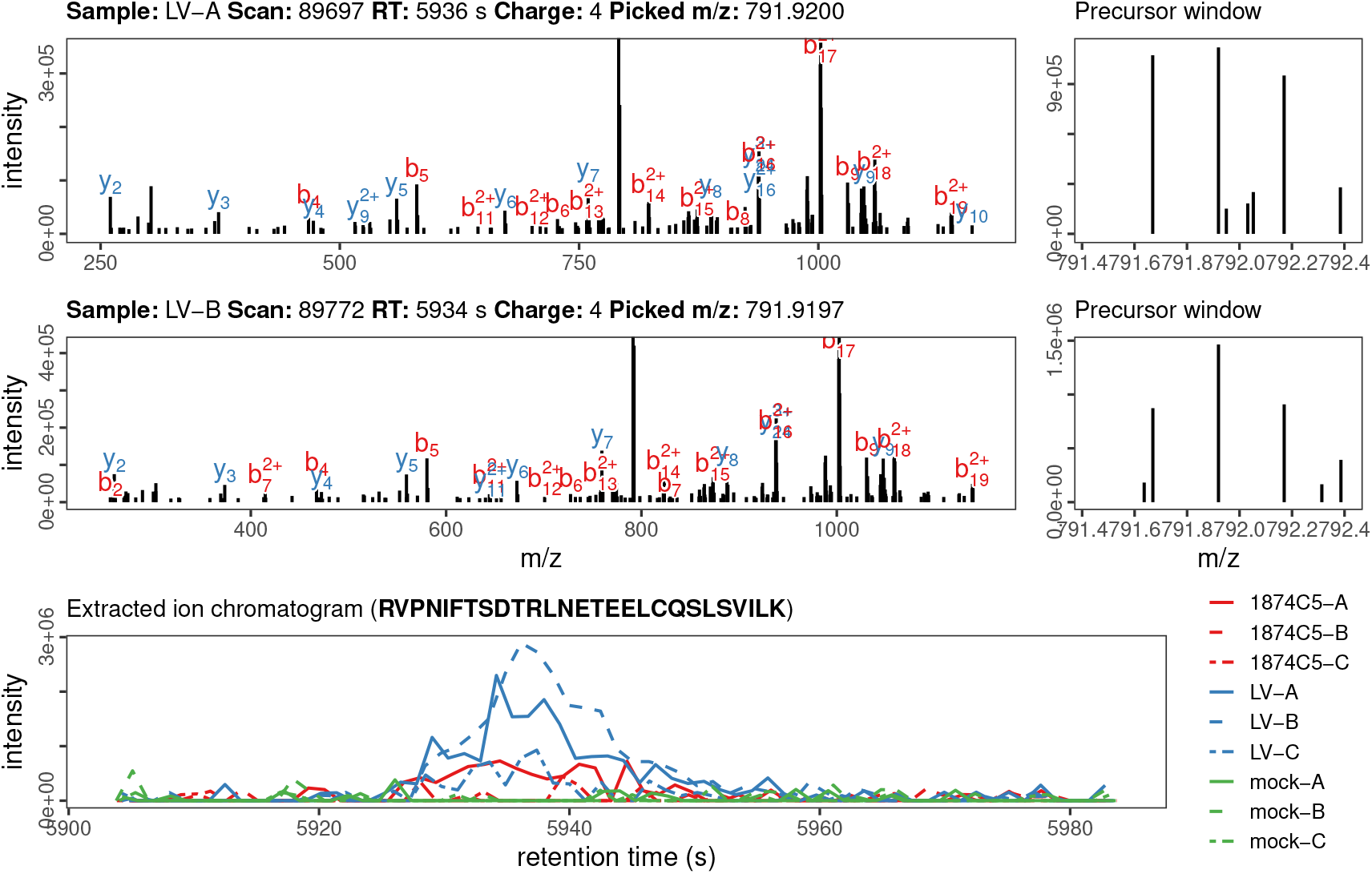
QC data for vIL-4 peptide RVPNIFTSDTRLNETEELCQSLSVILK identified by LC-MS/MS. Top rows show up to three MS2 spectra matched at 1% FDR, with matching y-b ion series labeled in blue and red. On the right of each row is the MS1 precursor window used for fragmentation. The bottom row shows an extracted ion chromatogram (XIC) for the peptide precursor mass over the matching retention time window for all nine samples (1875C5, LaryngoVac, and mock). True viral peptides should have XIC elution peaks in infected samples but not mock.

**Figure S16:**
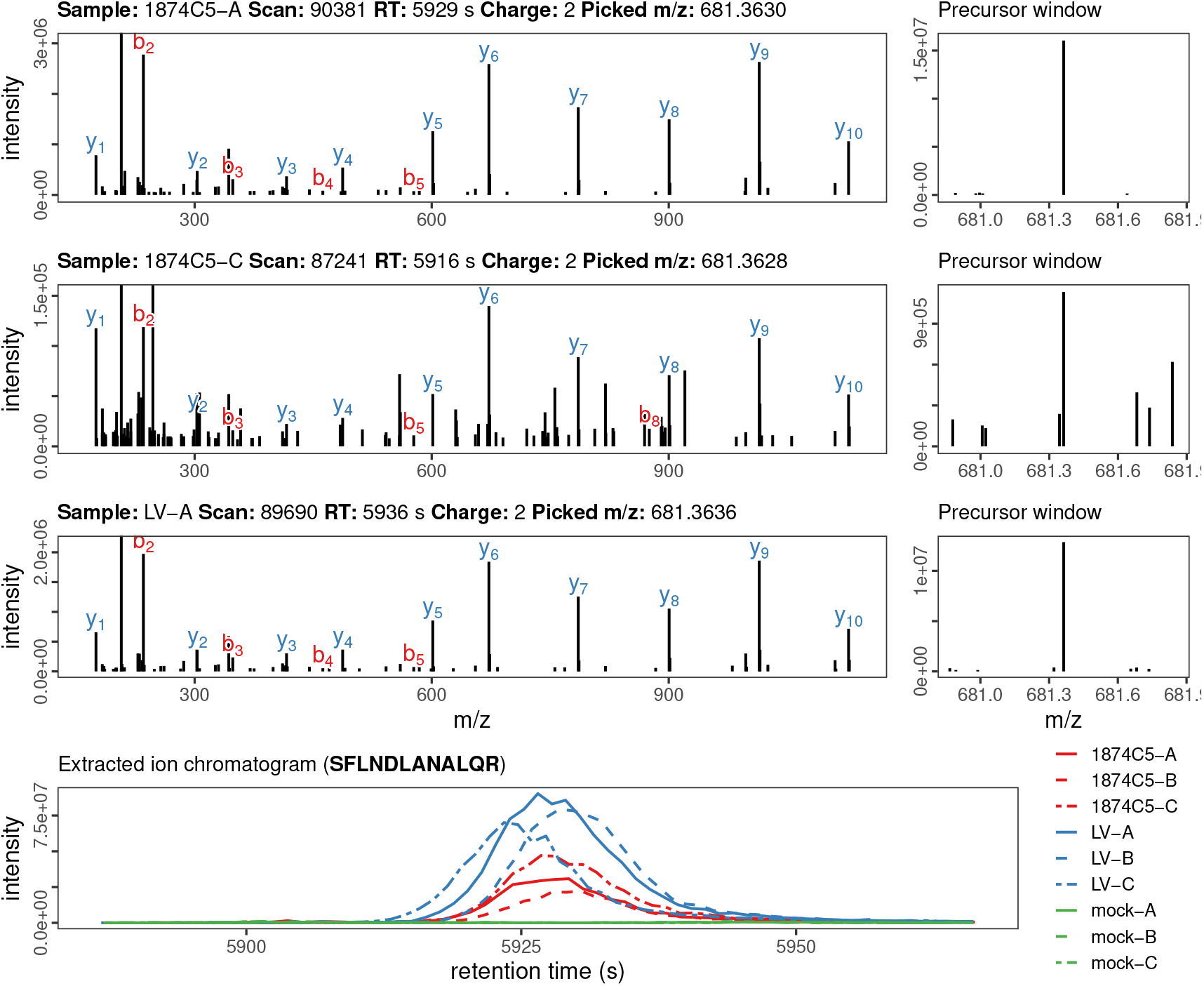
QC data for vIL-4 peptide SFLNDLANALQR identified by LC-MS/MS. Top rows show up to three MS2 spectra matched at 1% FDR, with matching y-b ion series labeled in blue and red. On the right of each row is the MS1 precursor window used for fragmentation. The bottom row shows an extracted ion chromatogram (XIC) for the peptide precursor mass over the matching retention time window for all nine samples (1875C5, LaryngoVac, and mock). True viral peptides should have XIC elution peaks in infected samples but not mock.

**Figure S17:**
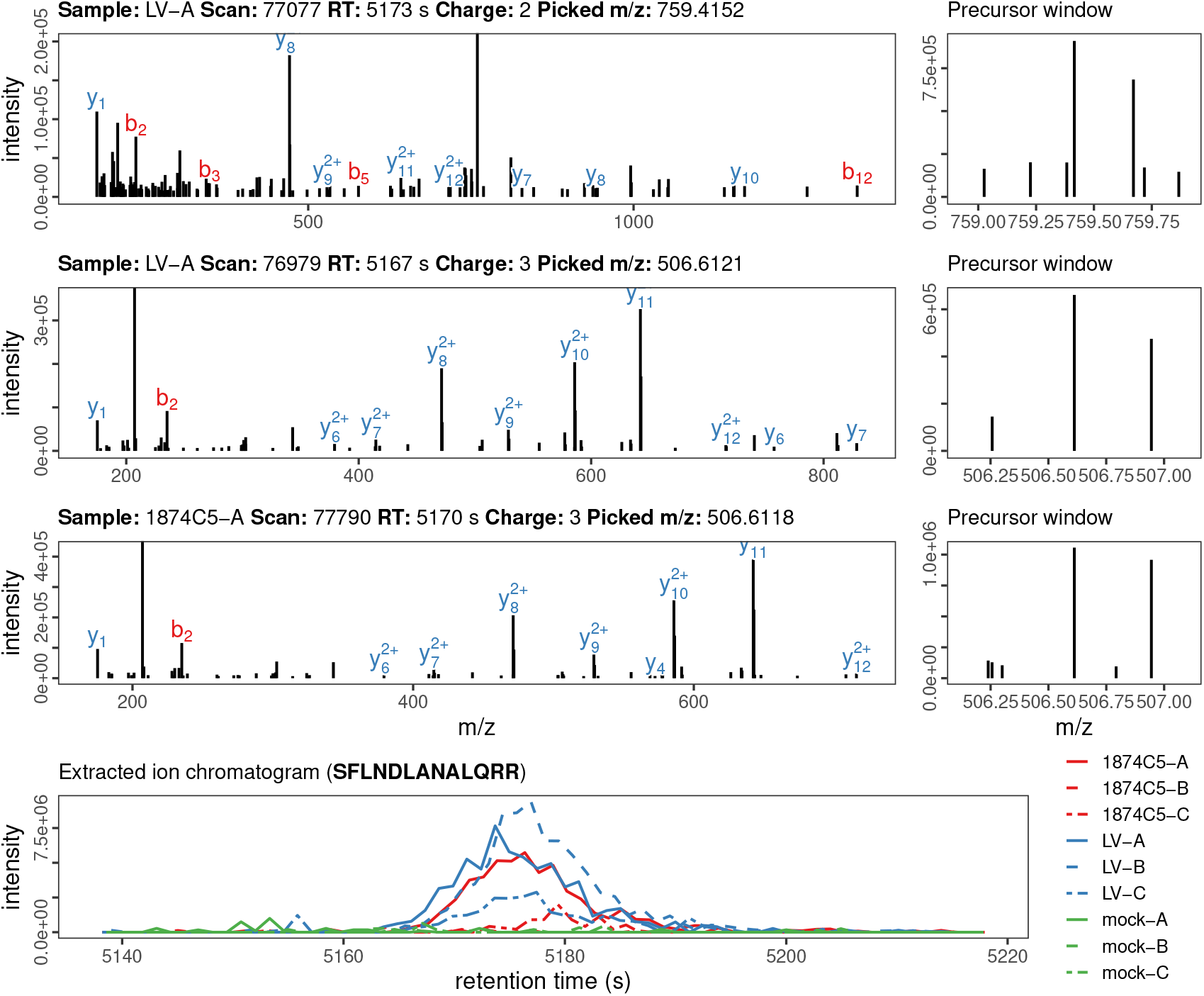
QC data for vIL-4 peptide SFLNDLANALQRR identified by LC-MS/MS. Top rows show up to three MS2 spectra matched at 1% FDR, with matching y-b ion series labeled in blue and red. On the right of each row is the MS1 precursor window used for fragmentation. The bottom row shows an extracted ion chromatogram (XIC) for the peptide precursor mass over the matching retention time window for all nine samples (1875C5, LaryngoVac, and mock). True viral peptides should have XIC elution peaks in infected samples but not mock.

**Figure S18:**
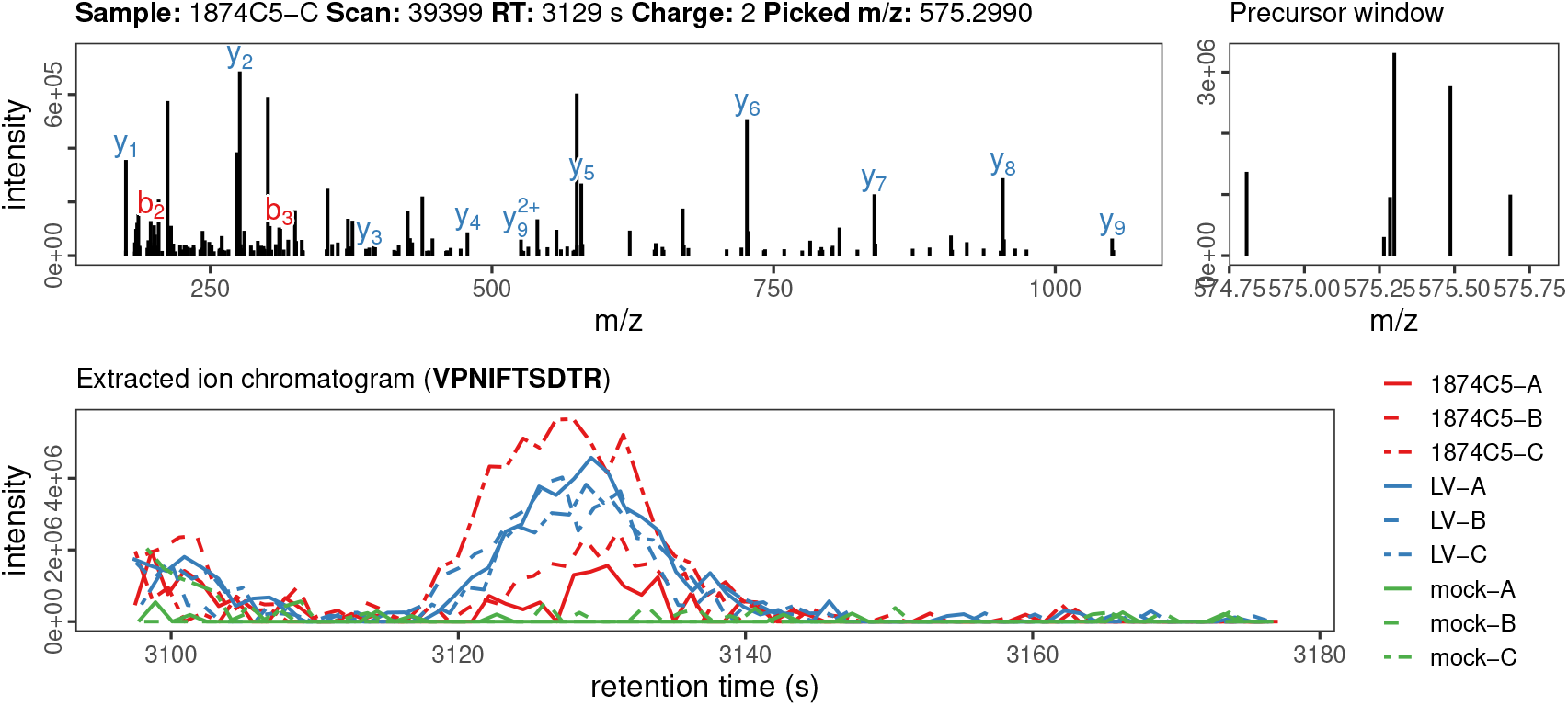
QC data for vIL-4 peptide VPNIFTSDTR identified by LC-MS/MS. Top rows show up to three MS2 spectra matched at 1% FDR, with matching y-b ion series labeled in blue and red. On the right of each row is the MS1 precursor window used for fragmentation. The bottom row shows an extracted ion chromatogram (XIC) for the peptide precursor mass over the matching retention time window for all nine samples (1875C5, LaryngoVac, and mock). True viral peptides should have XIC elution peaks in infected samples but not mock.

**Table S1:** Vertebrate IL-4 homologs used in the analysis. Accessions are from GenBank and are either the direct protein entries used or the genomic sequences on which the predicted genes are located (as indicated in the Notes).

| Common name | Scientific name | Accession | Notes |
| --- | --- | --- | --- |
| vIL-4 | Gallid alpha-herpesvirus 1 | N/A | from GaHV-1 1874C5 IsoSeq and internal 1874C5 corrected genome |
| chicken | Gallus gallus | NM_001398459.1 | used as-is |
| Chinese bamboo-partridge | Bambusicola thoracicus | POI22934.1 | miniprot/AUGUSTUS prediction, matches indicated protein accession |
| turkey | Meleagris gallopavo | ENSMGAT00000023360.1 | GenBank NM_001303181.1 is actually IL13 |
| ring-necked pheasant | Phasianus colchicus | NW_022205427.1 | miniprot/AUGUSTUS prediction |
| greater sage-grouse | Centrocercus urophasianus | JAHKSY010000026.1 | miniprot/AUGUSTUS prediction |
| rock ptarmigan | Lagopus muta | CM041489.1 | miniprot/AUGUSTUS prediction |
| green peafowl | Pavo muticus | JACDJE010000203.1 | miniprot/AUGUSTUS prediction |
| red-legged partridge | Alectoris rufa | CAMZON010000379.1 | miniprot/AUGUSTUS prediction |
| Japanese quail | Coturnix japonica | XM_015875546.1 | used as-is |
| guineafowl | Numida meleagris | XM_021410541.1 | used as-is |
| guan | Penelope pileata | WBMW01000786.1 | miniprot/AUGUSTUS prediction |
| bobwhite | Colinus virginianus | AWGT02000115.1 | miniprot/AUGUSTUS, structure differs from OXB76676 |
| brush-turkey | Alectura lathami | VXAV01004783.1 | miniprot/AUGUSTUS prediction |
| tufted duck | Aythya fuligula | XM_032196612.1 | used as-is |
| mallard | Anas platyrhynchos | NC_051785.1 | miniprot/AUGUSTUS, several nucleotide positions differ from MF346730.1 |
| black swan | Cygnus atratus | XM_035560814.1 | used as-is |

Table S1: Vertebrate IL-4 homologs used in the analysis. Accessions are from GenBank (*continued*)
| Common name | Scientific name | Accession | Notes |
| --- | --- | --- | --- |
| egret | Egretta garzetta | XM_009647167.1 | used as-is |
| Adelie penguin | Pygoscelis adeliae | XM_009322783.1 | used as-is |
| goshawk | Accipiter gentilis | XM_049829109.1 | used as-is |
| golden eagle | Aquila chrysaetos | XM_029998407.1 | used as-is |
| zebra finch | Taeniopygia guttata | NC_044225.2 | miniprot/AUGUSTUS, structure differs from XM_041718987.1 |
| song sparrow | Melospiza melodia maxima | RZID01002771.1 | used as-is |
| ostrich | Struthio camelus | JJRT01005166.1 | miniprot/AUGUSTUS, structure differs from XM_009679367.1 |
| emu | Dromaius novaehollandiae | NW_020453704.1 | miniprot/AUGUSTUS, structure differs from XP_025974110.1 |
| tinamou | Nothoprocta perdicaria | NW_020454172.1 | miniprot/AUGUSTUS, structure differs from XM_026045620.1 |
| saltwater crocodile | Crocodylus porosus | NW_017728914.1 | miniprot/AUGUSTUS prediction |
| alligator | Alligator sinensis | XM_006023567.1 | used as-is |
| slider turtle | Trachemys scripta elegans | NC_048305.1 | miniprot/AUGUSTUS, structure differs from XM_03477890.1 |
| green sea turtle | Chelonia mydas | NC_057854.1 | miniprot/AUGUSTUS prediction |

**Table S2:**
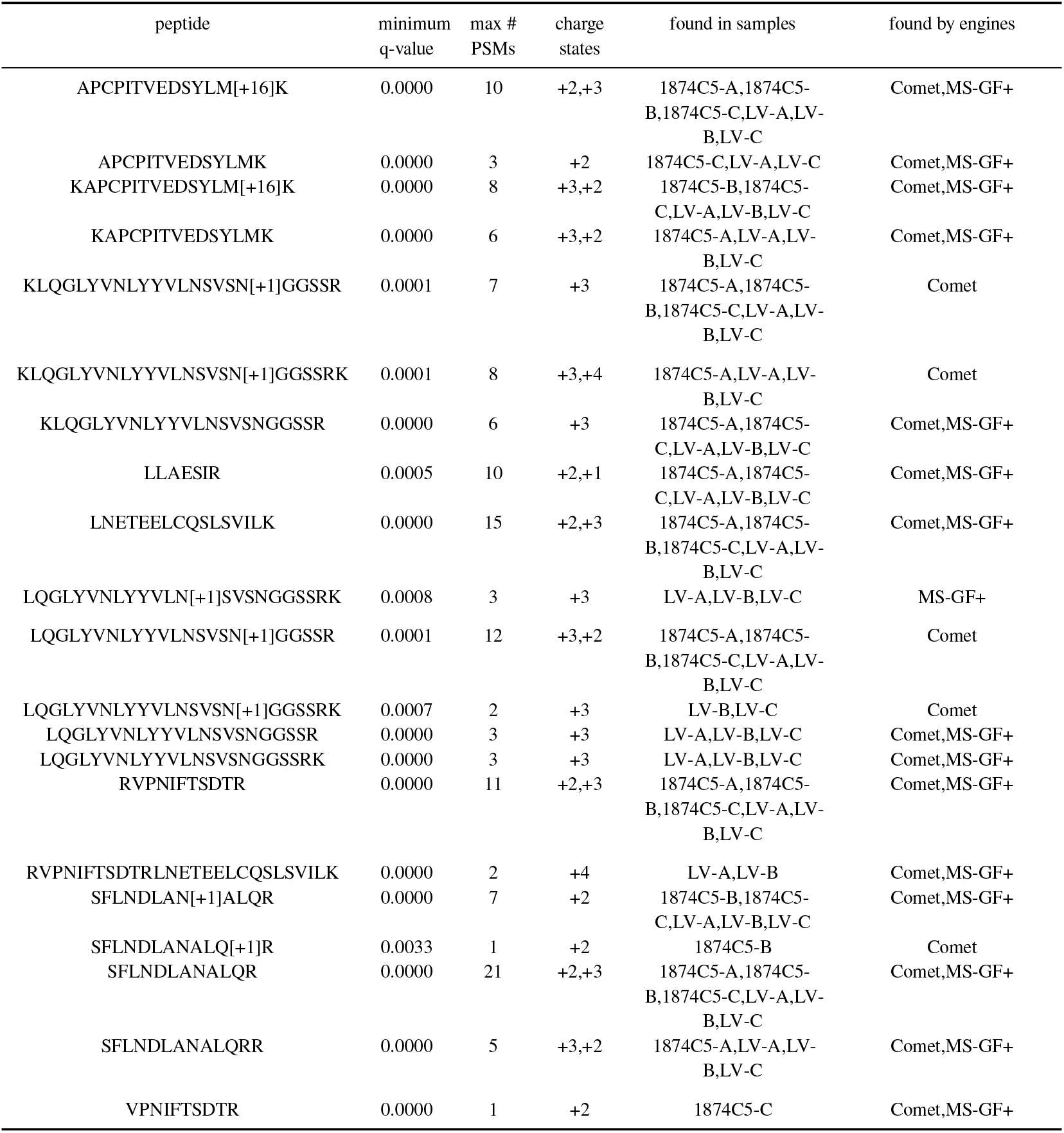
All vIL-4 peptide proteoforms detected by LC-MS/MS of infected cell culture at an FDR of 1%. Columns shown are: peptide sequence with post-translation modification (PTM) masses in square brackets; minimum q-value assigned (from all samples); maximum per-sample PSM count observed; observed charge states across all samples; replicates the peptide proteoform was detected in; search engines identifying the specific proteoform. Note that where PTMs could not be confidently localized to a single amino acid, different search engines may prefer and report different modified positions. An example is the deamidated form of peptide LQGLYVN-LYYVLNSVSNGGSSRK.

